# Lewy pathology in Parkinson’s disease consists of a crowded organellar, membranous medley

**DOI:** 10.1101/137976

**Authors:** Sarah H. Shahmoradian, Amanda J. Lewis, Christel Genoud, Jürgen Hench, Tim Moors, Paula P. Navarro, Daniel Castaño-Díez, Gabriel Schweighauser, Alexandra Graff-Meyer, Kenneth N. Goldie, Rosmarie Sütterlin, Evelien Huisman, Angela Ingrassia, Yvonne de Gier, Annemieke J.M. Rozemuller, Jing Wang, Anne De Paepe, Johannes Erny, Andreas Staempfli, Joerg Hoernschemeyer, Frederik Großerüschkamp, Daniel Niedieker, Samir F. El-Mashtoly, Marialuisa Quadri, Wilfred F.J. van IJcken, Vincenzo Bonifati, Klaus Gerwert, Bernd Bohrmann, Stephan Frank, Markus Britschgi, Henning Stahlberg, Wilma D. J. van de Berg, Matthias E. Lauer

## Abstract

Parkinson’s disease, the most common age-related movement disorder, is a progressive neurodegenerative disease with unclear etiology. Key neuropathological hallmarks are Lewy bodies and Lewy neurites, which are neuronal inclusions that are immunopositive for the protein α-synuclein. In-depth ultrastructural analysis of this Lewy pathology is crucial to understanding pathogenesis and progression of the disease. Using correlative light and electron microscopy/tomography on brain tissue from five Parkinson’s disease brain donors, we identified α-synuclein immunopositive Lewy pathology and could show that the majority of these features including Lewy bodies and Lewy neurites primarily consists of a crowded membranous medley of vesicular structures and dysmorphic organelles. Only a small fraction of observed Lewy bodies contained predominant proteinaceous filaments, as previously described. The crowding of organellar components was confirmed by STED-based super-resolution microscopy, and high lipid content within the α-synuclein immunopositive inclusions was corroborated by confocal imaging, CARS/FTIR imaging and lipidomics. Applying this correlative high-resolution imaging and biophysical approach, we discovered in the postmortem brain of Parkinson’s patients a subcellular protein-lipid compartmentalization not previously described in Lewy pathology.

## Introduction

Lewy bodies (LB) have been recognized as the main pathological hallmark of Parkinson’s Disease (PD) and Dementia with Lewy Bodies (DLB) following their microscopic discovery over 100 years ago. The protein α-synuclein (aSyn) has been found to be a major component of Lewy bodies and is considered to play a central role in their formation and of other Lewy pathologies including Lewy bodies, pale bodies, and Lewy neurites^1^. The morphology of LB revealed by light microscopy (LM) varies depending on their location in the brain (brainstem, limbic or neocortical)^2, 3^, may reflect maturation stage and genetic background, and their biochemical and proteomic composition is known to be complex^2, 4^. The morphology of Lewy neurites, displaying the same aSyn immunohistochemical staining profile^5, 6^ as those found in the neuronal cell bodies, also varies between brainstem and (sub)cortical brain regions^7^.

The processes by which Lewy pathology arise and their role in neurodegeneration remain elusive. The current, leading hypothesis in PD research proposes that intraneuronal aSyn first forms as abnormal oligomers, possibly induced by extracellular pathogenic aSyn aggregates that were taken up and subsequently transforms into β-sheet rich amyloid fibrils, which are the basis of the LB^8^. Transmission electron microscopy (TEM) carried out on sarcosyl-insoluble fractions of brain tissue extracts or formalin-fixed, de-paraffinized and resin-embedded postmortem brain tissue of patients with PD or DLB, revealed the ultrastructure of LB as filaments immunoreactive for aSyn^3, 9^. By contrast, evidence from ultrastructural studies based on neuroanatomical localization point to an electron-dense core and granular features^10^. Proteome studies have shown that LB consists of more than 300 proteins, of which approximately 90 have been confirmed by immunohistochemistry in various postmortem studies and are associated with aSyn, protein degradation systems, molecular chaperones or axonal damage^2^. Current literature about the nature of Lewy pathology is in several aspects still incomplete and requires validation in order to make proper conclusions about pathogenesis and progression of Parkinson’s disease. A clearer understanding of the building blocks of Lewy pathology is therefore urgently needed.

The discovery that recombinant aSyn can form filaments *in vitro* has most likely influenced the search for a filamentous type of aSyn-immunopositive inclusion within human brain tissue and extracts in past ultrastructural studies. Alarmingly, to date, there exists no ultrastructural study using postmortem brain tissue from multiple PD donors that used unequivocal identification of Lewy pathology from an unbiased correlative microscopy approach. With the advent of modern technologies for electron microscopy such as energy filters, direct electron detectors, and drift-correcting software for tomography, we now have the possibility to obtain a clearer and more accurate picture of the 3D structure of such aSyn-immunopositive pathological inclusions, including the capability of distinguishing amyloid fibrils from lipid membranes. Importantly, such advanced transmission electron microscopy imaging can be done in correlation with light microscopy imaging. Here, we present an unprecedented 3D view of the structural components of LB and LN in well-preserved brain tissue from five PD brain donors using correlative light and electron microscopy (CLEM). The correlative methods employed clearly show that the vast majority of Lewy pathology is comprised of crowded membranous material rather than predominantly filamentous structures, and suggest that the membranes originate from vesicles and fragmented organelles, including mitochondria. Serial block-face scanning electron microscopy (SBFSEM)^11^, an EM method which enables visualizing a larger tissue volume at intermediate resolution as compared to CLEM, also revealed crowded organelles and membrane fragments in cellular aggregates in the SN, and a shell of mitochondria surrounding some of these inclusions. Correlative multi-labeling and stimulated emission depletion microscopy (STED) were employed to corroborate these assignments in brain tissue from 14 PD donors, including those used for CLEM, and demonstrated the crowding of aSyn, lipids, lysosomal structures and mitochondria in LB. The presence of lipids in LB was confirmed by label-free compositional mapping methods and mass spectrometry. The nanoscale information obtained suggests that the membrane crowding observed in Lewy pathology is modulated by aSyn, and supports the hypothesis that impaired organellar trafficking contributes to PD pathogenesis^12^.

## Results

A correlated light and electron microscopy (CLEM) approach was employed to identify Lewy pathology in advanced PD brain donors (Table 1). By CLEM, we obtained a 3D view of the ultrastructure of 16 aSyn-immunopositive inclusions in neurons (Supplementary Table 1), three of which were identified as Lewy neurites (LN), within postmortem samples from five PD donors (Table 1). The presence, distribution and morphology of Lewy pathology in the *substantia nigra* SN and hippocampal CA2 in these donors were determined by light microscopy (LM) of paraffin-embedded tissue sections immunostained for aSyn and illustrated the accumulation of aSyn in these brain regions (Fig. S1). CLEM (Fig. S2-S12) was performed using adjacent tissue blocks prepared at autopsy in parallel from the same tissue sample (herein referred to as parallel tissue blocks); aSyn immunoreactive inclusions were identified by histological staining followed by LM, and 3D transmission electron microscopy (TEM) tomograms (Figs. 1-3, S5; Movies 1-19) and 2D TEM images (Figs. 4, S7-S12) were recorded from adjacent tissue sections (*i.e.*, regions maximally 450 nm above or below in the block) by TEM. Of all Lewy pathology that we identified using CLEM, we observed only a single aSyn-immunopositive inclusion amongst neuromelanin-containing organelles in the *substantia nigra* of Donor C-PD comprised of a proteinaceous core, radiating filaments and organelles at the periphery (Fig. 1d, S8 and Movie 4, Supplementary Table 1). Also within neuromelanin-containing organelles in the *substantia nigra* of two separate brain donors, we observed strongly aSyn-immunopositive inclusions containing abundant aggregates of mitochondria, numerous lipid vesicles and worm-like tubulovesicular structures (Fig. S9: Donor D-PD, Fig. S10: Donor E-PD, pink arrowheads), interspersed with randomly oriented filaments (Figs. S9-S10, blue arrowheads). In two aSyn-immunopositive inclusions not visible with neuromelanin yet also found in the *substantia nigra* of Donors D-PD and E-PD, we observed both abundant vesicular structures and structures with filamentous appearance. In Donor D-PD, we observed abundant tubulovesicular structures (Fig. S11, pink arrowheads) interspersed with randomly oriented filaments (Fig. S11, blue arrowheads). In Donor E-PD, we observed a single aSyn-immunopositive inclusion comprised of autophagic vacuolar-like structures (membrane-enclosed, “empty” vesicles) similar to what was observed in a LN (Fig. 4), vesicles with a ruffled border, distorted vesicles, vesiculotubular structures (Fig. S12, pink arrowheads), and randomly oriented filaments (Fig. S12, blue arrowheads). Only three aSyn-immunopositive inclusions amongst 16, appeared to consist primarily of filamentous structures (Fig. 1d, S6d-e). Overall, the majority of aSyn inclusions appear to mainly consist of a crowded organellar and membranous medley (Figs. 1a-c, 2-6, S6a-c, e-f, and Movies 1-3 and 5-7, 9-17, 20-21). The TEM tomograms revealed mitochondria and numerous cellular organelles clearly visible at the periphery and interior of the inclusions. Densely compacted membranous structures, tubulovesicular structures, distorted vesicles and haphazardly distributed filaments were also visible.

**Figure 1.**
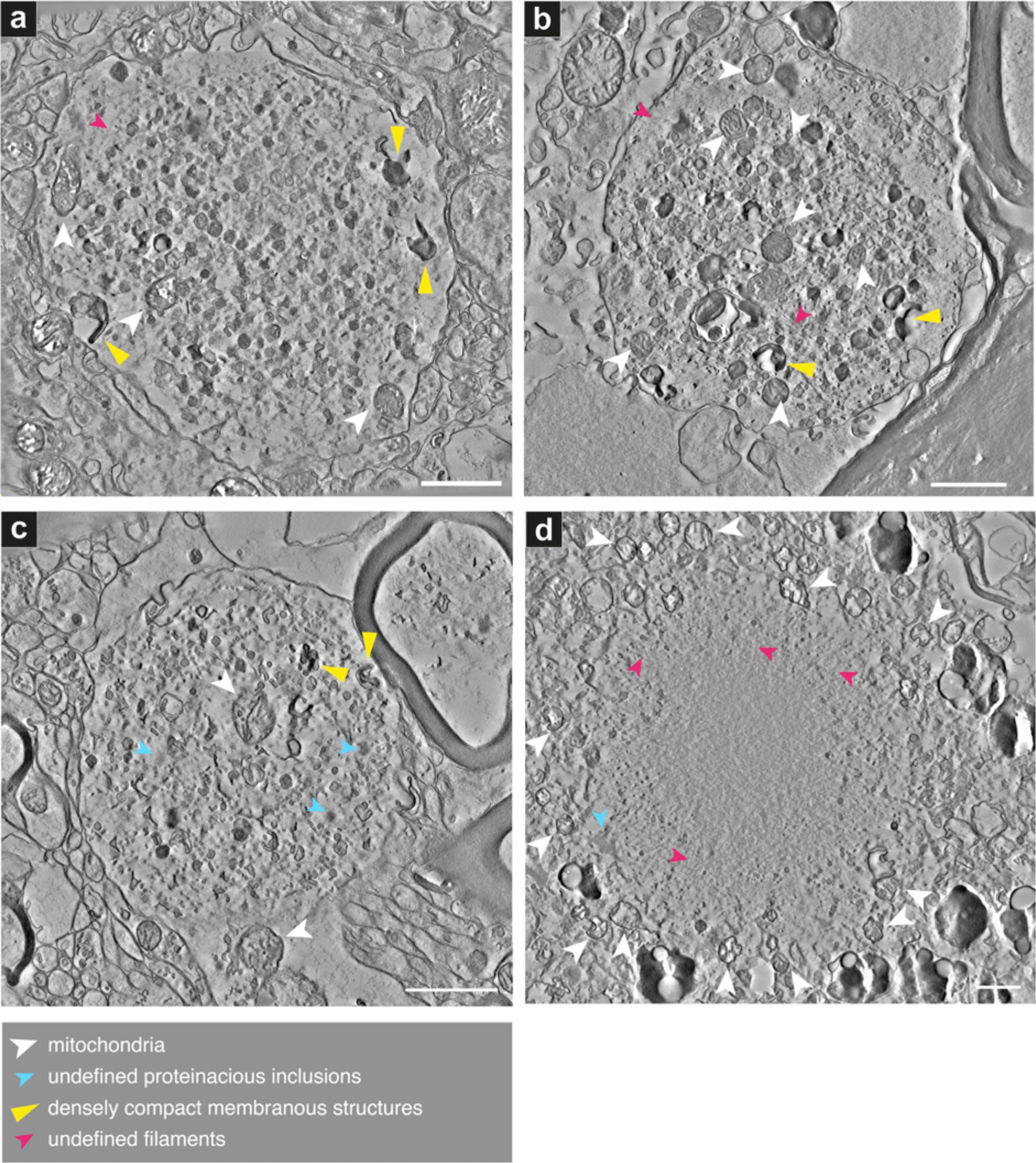
Lewy pathology shows abundant membranous structures, abnormal organelles and vesicles. Projections of the central 20 slices of each reconstructed 3D tomogram are shown for each aSyn-immunopositive inclusion found by CLEM, and surrounding cellular milieu. Feature details (arrowheads) are tabulated in Supplementary Table 1. Additional aSyn-immunopositive Lewy pathological inclusions are shown in Figs. 3, 4 and S5-S12. Donor identities are shown in Table 1. (a) Donor A-PD (Movie 1), (b) Donor B-PD (Movie 2), (c) Donor D-PD (Movie 3), (d) Donor C-PD (Movie 4), CLEM data shown in Fig S3a. Scale bars = 1 µm.

**Figure 2.**
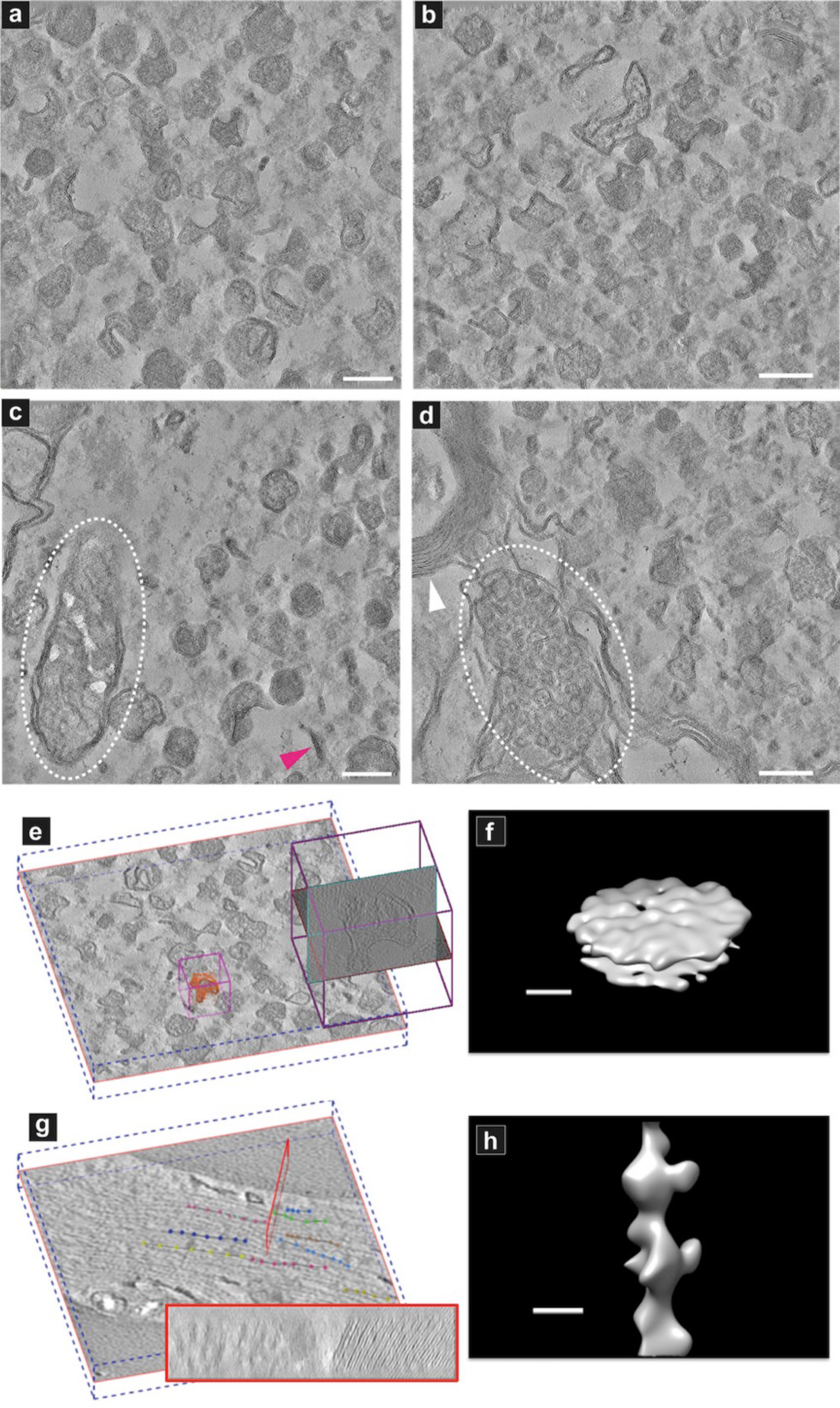
Electron tomography and subtomogram averaging reveal membranous nature of Lewy pathology. Projections of the central 60 slices of each reconstructed 3D tomogram are shown for each sub-region of aSyn-immunopositive inclusion shown in a-d for Donor A-PD. (a) Inner region of the inclusion as shown in Fig. 1a and (b) in Fig. S6a. (c) Edge of the inclusion as shown in Fig. 1a. White oval = distorted mitochondrion, pink arrow = representative disc-like membranous structure; more of such structures are visible in 3D, Movies 11-15. (d) Edge of the inclusion as shown in Fig. S6a. Cluster of vesicular structures in adjacent yet separate compartment to the inclusion is visible (white oval). (e) Z-orthoslice in a tomogram from an inclusion and the region selected for sub-tomogram averaging (pink box). Subvolumes were sampled on high intensity points around the indicated locations (red). (f) Subtomogram average for the sampled densities in the inclusion revealed a membrane structure with the two leaflets separated by the typical spacing of lipid bilayer. (g) Z-orthoslice in a tomogram from a neurite of a non-neurological, age-matched control brain donor. The red box indicates a cross-section showing myelin sheaths (right side of box) and filaments (left side of box) within the neurite. (h) Subtomogram average for the sampled densities along the dotted lines in ‘g’ revealed a filamentous structural signature. Scale bars a-d = 200 nm; f, h = 10 nm.

**Figure 3.**
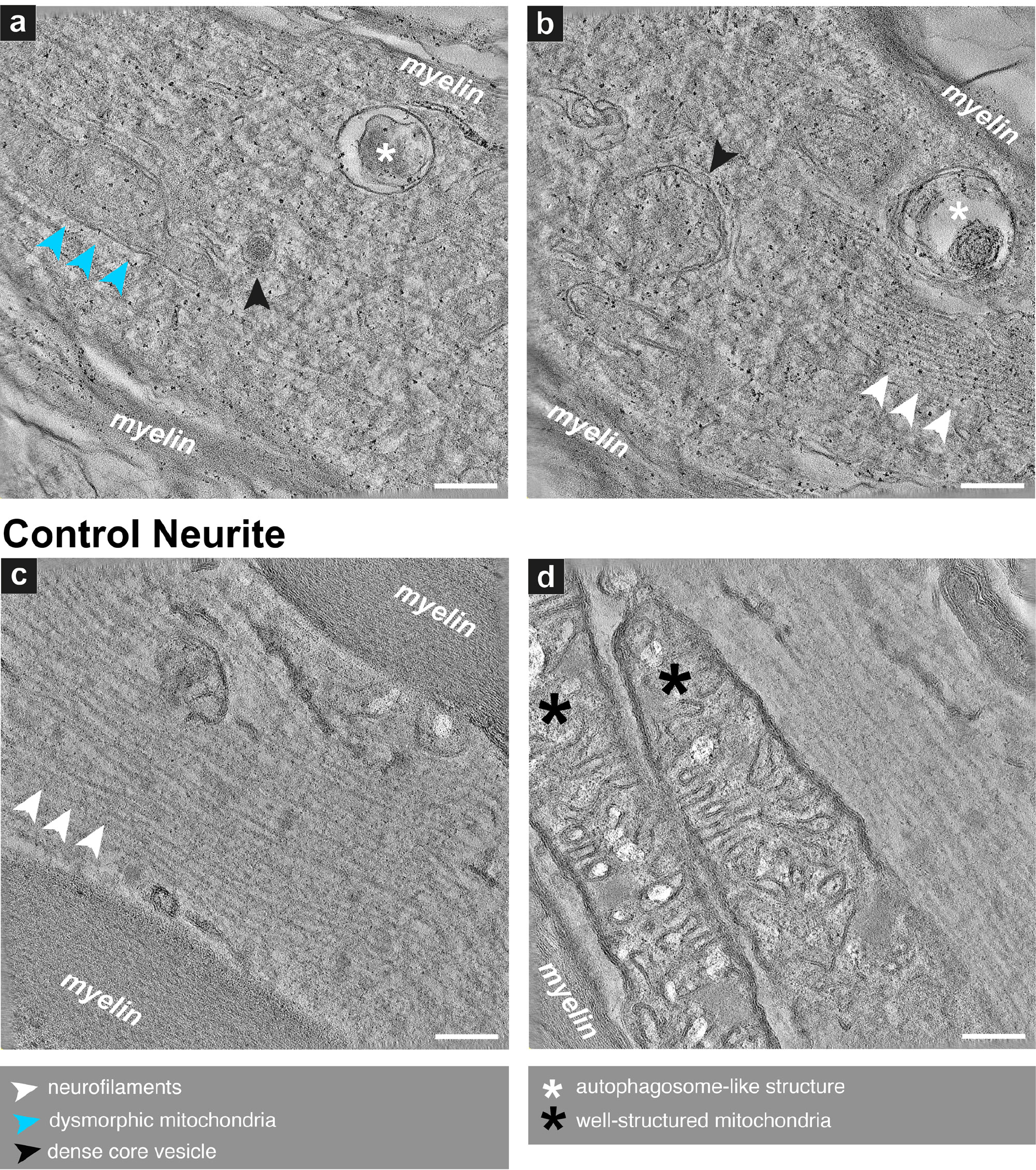
Nigral Lewy neurite reveals disrupted cytoskeletal elements, dysmorphic mitochondria and autophagosome-like structures. Electron tomogography of LN identified using CLEM from Donor B-PD *substantia nigra*. (a) Inner region of the LN shown in the overview image Fig. S7 (upper dotted red box) and Movie 16. Elongated double-membrane enclosed structures with faint inner membrane convolutions are visible, identified as abnormally elongated and otherwise dysmorphic mitochondria (blue arrowheads). (b) Different inner region of the LN shown in Fig. S7 (lower dotted red box) and Movie 17. **Control neurites**: (c) Inner region of a normal neurite (Movie 18) in the brain of a non-neurological, age-matched control donor (Donor F-Control, Table 1). Ordered neurofilaments are visible, flanked by a well-structured myelin sheath representing the enclosing axon. (d) Inner region of a separate neurite (Movie 19) in the brain of the same donor as shown in ‘c’. a, b = Projection through the central 40 slices of the reconstructed 3D tomogram; c, d = Projection through the central 60 slices of the reconstructed 3D tomogram. Scale bars = 200 nm.

**Figure 4.**
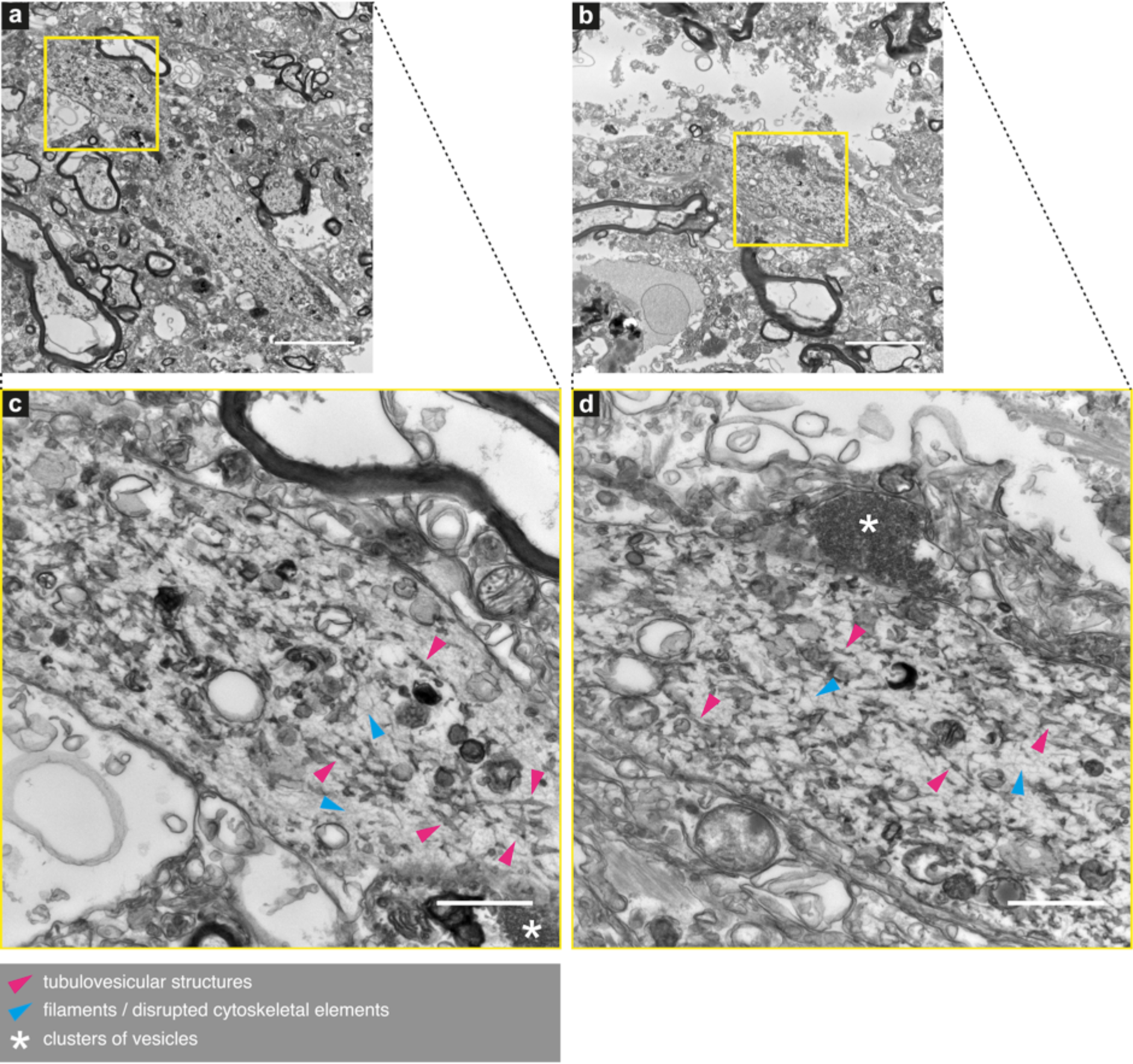
Nigral Lewy neurite revealing disrupted cytoskeletal elements, tubulovesicular, lysosome- and autophagosome-like structures. Electron tomography of a LN identified by CLEM from Donor E-PD *substantia nigra*. 2D images. Top part (a) and bottom part (b) of the LN as shown in Fig. S5. (c, d) Inner region at higher magnification of the LN as shown in ‘a’ and ‘b’, respectively. Autophagic vacuolar-like structures (membrane-enclosed, “empty” vesicles) and lysosomal structures such as lipofuscin (black/dark gray semi-circular structures) and structures resembling mitochondria are visible in addition to the annotated features. Scale bars: a, b = 5 µm, c, d = 1 µm.

**Figure 5.**
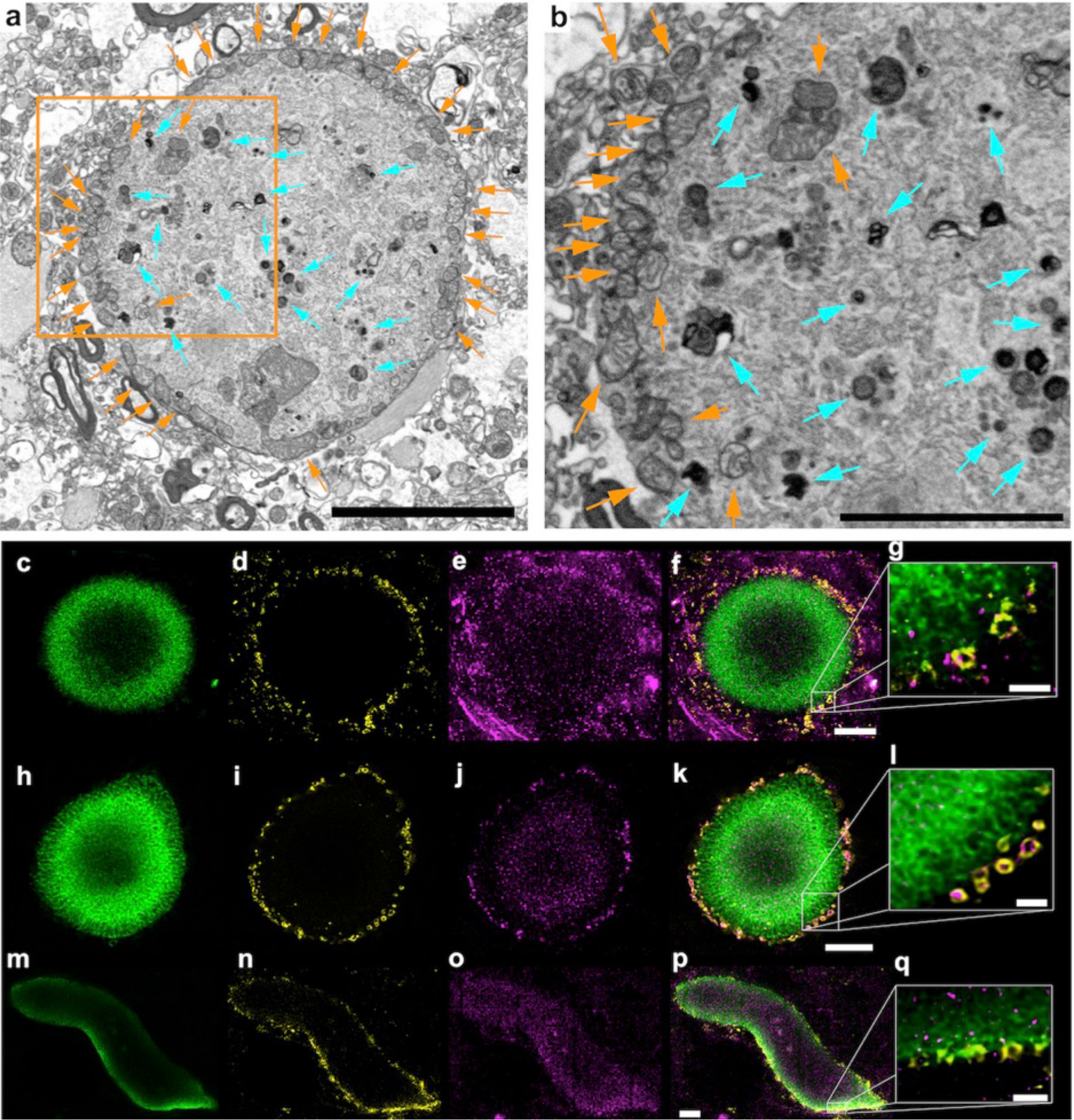
Sub-cellular features of Lewy pathology reveal the organelle distribution. (a) SBFSEM imaging of heavy-metal stained *substantia nigra* (SN) tissue in Donor B-PD showing a surrounding ring of mitochondria (orange arrows) and structures reminiscent of lysosomes (aqua arrows) further within the inclusion. (b) Enlarged view of the boxed region in ‘a’, similarly annotated. (c-q) Microscopy of separate inclusion in the same SN region of the same brain donor (Donor B-PD) (c-g), and Lewy pathology in the SN of Donor A-PD (h-q), showing the distribution of (c, h, m) marker for phosphorylated aSyn (pS129), (c, i, n) marker for mitochondria (porin/VDAC1), (e, j, o) marker for lysosomes (LAMP1), (f, k, p) overlay of all markers, (g, l, q) enlarged view of the edge of the Lewy pathology in ‘f’, ‘n’, and the Lewy neurite in ‘p’. Images are representative across 14 PD donors for Lewy pathological inclusions with an outer layer of p-aSyn. Scale bars: f, k, p = 5 µm; g, l, q = 1 µm.

**Figure 6.**
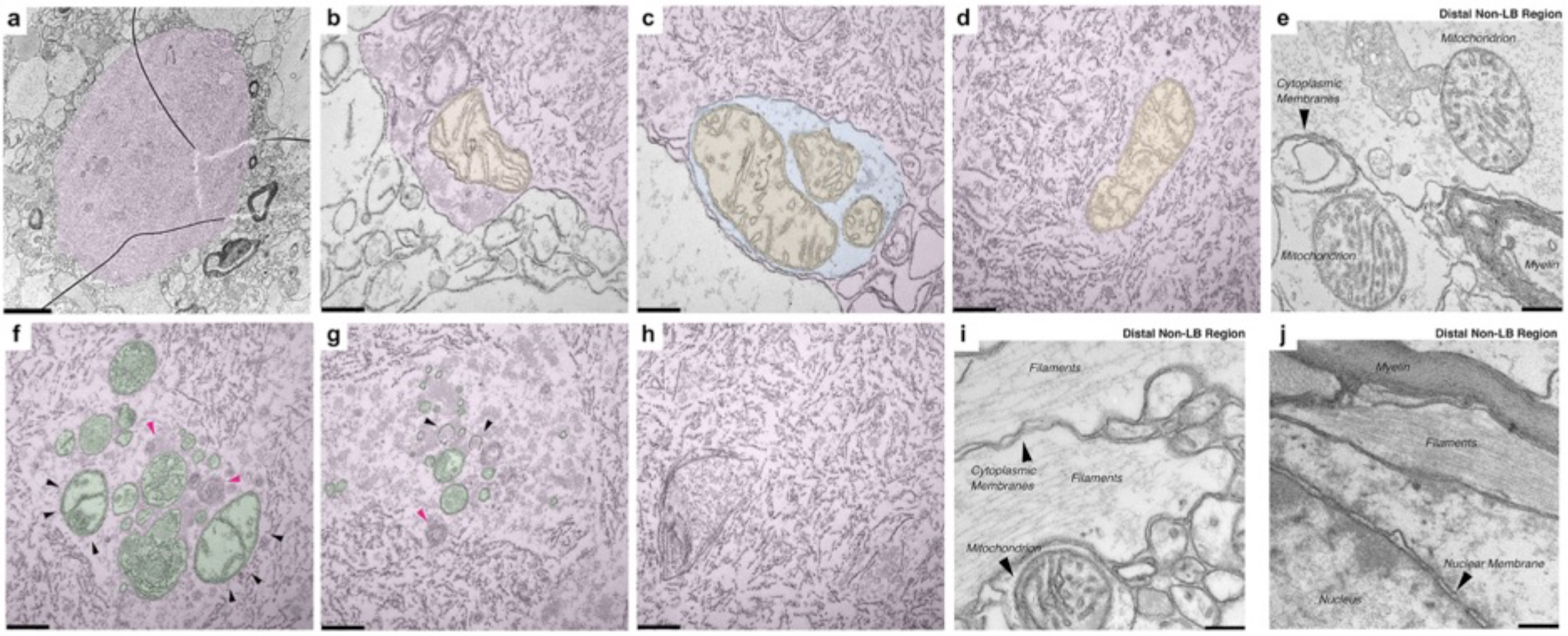
Inner architecture of Lewy pathology analyzed by correlative SBFSEM/TEM shows membrane fragments and organelles. TEM images of sections (50 nm thick) of an aggregate localized by SBFSEM in the SN of Donor B-PD. Images are artificially colored: overall inclusion = pink, membrane compartments = blue, mitochondria-orange, other organelles = green. **(a)** Overview of the Lewy pathological inclusion within the tissue. **(b)** Edge of the inclusion with a mitochondrion present. **(c)** Another edge of the inclusion with mitochondria enclosed in a membrane-delimited compartment. **(d)** Mitochondrion surrounded by membrane fragments within the inclusion. **(e)** Two apparently normal mitochondria with clear cristae in an area within the same tissue section but distal to the region with the inclusion. Cytoplasmic membranes and a myelin sheath are also present. **(f)** Putative mitochondria (black arrowheads) and putative lysosomes (pink arrowheads) within the inclusion. **(g)** Membrane-bound structures, some potentially omegasomes (black arrowheads) and one a putative lysosome (pink arrowhead), surrounded by fuzzy structures that may represent proteinaceousnacious deposits within the inclusion. **(h)** General cluster of membrane fragments appearing within the inclusion. **(i)** Typical cytoskeletal filaments along with mitochondrion showing clear cristae, in an area within the same tissue section but distal to the region with the inclusion. **(j)** Condensed filaments and myelin along with nuclear membrane, in an area within the same tissue section but distal to the region with the inclusion. Note the appearance of membranes vs. filamentous structures; the staining approach employed made it easy to distinguish between them. Scale bars: a = 1.5 µm, all others = 200 nm.

**Table 1.** Clinical and pathological characteristics of brain donors. Donors are referred in the text as either ‘PD’ or ‘Control’ in the following nomenclature: Donor A-PD, Donor E-Control, etc. Staging of Lewy body disease and Alzheimer’s disease related pathology was performed based on Brain Net Europe (BNE) consensus guidelines (aSyn^66^; NFT^67^; Abeta^68^) and National Institute on Aging-Alzheimer’s Association (NIA-AA) criteria^69^. Additionally, brain tissue was also inspected for other salient pathology such as age-related tau astrogliopathy (ARTAG)^70^, cerebral white matter rarefactions, cerebral amyloid angiopathy (CAA^71^) and (micro)infarctions and hemorrhages and TDP pathology^72^. Load of alpha-synuclein pathology was high in limbic and neocortical regions in all PD(D) cases (Braak aSyn stage 5-6), and absent (0) in the control (“Con”) case. Neurofibrillary and amyloid-β (Abeta) pathology were absent (0) or low (1-2) in most PD(D) and control, but two cases showed more severe NFT (3) or Abeta) load (C). Eight cases showed mild (1) or severe (2) capillary cerebral amyloid angiopathy (CAA). None of the cases showed infarcts or TDP pathology. PDD = Parkinson’s disease with dementia; age-at-onset = age at clinical diagnosis of PD; n.a. = not available; Con = non-neurological control; NFT = neurofibrillary tangles; CERAD = Consortium to Establish a Registry for Alzheimer Disease; CAA = cerebral amyloid angiopathy; aSyn = α-synuclein; PMD = postmortem delay. Age at onset = age at clinical diagnosis of PD. * = Donors used in EM studies. All donors were used for STED studies.

| Donor | Diagnosis | Age at onset | Age at death | Sex | Brain weight (g) | PMD (hrs: min) | Braak NFT stage | CERAD (Abeta) score | Braak aSyn stage | Thal phase | CAA type |
| --- | --- | --- | --- | --- | --- | --- | --- | --- | --- | --- | --- |
| A* | PDD | 59 | 77 | M | 1240 | 5:15 | 2 | 0 | 6 | 3 | 2 |
| B* | PDD | 75 | 90 | F | 1335 | 4:45 | 3 | B | 6 | 3 | 2 |
| C* | PD | 76 | 84 | M | 1430 | 4:50 | 3 | 0 | 5 | 0 | 0 |
| D* | PDD | 66 | 74 | M | 1350 | 5:15 | 2 | 0 | 6 | 3 | 1 |
| E* | PD | n.a. | 63 | M | n.a. | 5:55 | 2 | 0 | 5 | 0 | 0 |
| F* | Con | n.a. | 72 | F | 1205 | 5:15 | 1 | A | 0 | 1 | 1 |
| G | PD | 55 | 69 | M | 1405 | 6:50 | 1 | O | 6 | 0 | T2 |
| H | PD | 59 | 83 | M | 1335 | 5:15 | 1 | O | 6 | 1 | no |
| I | PDD | 67 | 80 | M | 1279 | 5:30 | 2 | B | 6 | 2 | T2 |
| J | PDD | 64 | 72 | M | 1210 | 4:00 | 1 | B | 6 | 3 | no |
| K | PD | 61 | 76 | F | 1248 | 5:30 | 1 | O | 6 | 1 | no |
| L | PDD | 69 | 80 | M | 1180 | 5:25 | 1 | O | 6 | 1 | no |
| M | PDD | 72 | 87 | F | 1135 | 5:25 | 2 | A | 6 | 2 | no |
| N | PDD | 78 | 83 | M | 1256 | 5:15 | 1 | B | 6 | 3 | T2 |
| O | PDD | 67 | 80 | F | 1218 | 5:15 | 1 | C | 6 | 3 | T2 |

Besides abundant vesicular structures scattered throughout the aSyn-immunopositive inclusions, dense “dark” L-shaped structures resembling stacks of compacted membrane sheets with a membrane layer spacing of ∼6 nm were distinguished (Fig. 1a-c, S6a-c, e, yellow arrowheads; Movies 1-3, 5-7, 9); herein referred to as tailed membrane stacks. They cannot be attributed to myelin, since the average membrane spacing of myelin sheaths is 10.7 nm in the central nervous system^13^. Furthermore, vesicular structures of varying electron density, some filled with more material than others (Fig. 2a-d; Movies 11-15), were clearly observed within the inclusions in higher magnification TEM tomograms. These structures are reminiscent of lysosomes and autophagosomes^14^. In addition, the tomograms reveal distorted mitochondria (Figs. 1 and S6, white arrowheads, 2c dotted ellipse; Fig. 3b black arrowhead, Fig. 6b-d, Figs. S9-S10) and other features (Fig. 2c pink arrowhead) that resemble the disk-like structures reported for *in vitro*-generated aSyn-lipoprotein particles examined by cryo-EM^15^. These disk-like and tubulovesicular structures appear frequently in 2D TEM images of other aSyn-immunopositive structures in different brain donors (Figs. S9-S12, pink arrowheads) and within two regions of a LN in Donor E-PD (Fig. 4, pink arrowheads). Computational analysis of the vesicular structures (Fig. 2e) by subtomogram averaging revealed their 3D structure, indicating the presence of two basal planes as expected for membrane leaflets (Fig. 2f). As a control for the subtomogram averaging, neurofilaments present within a neurite from a non-neurological control human brain (Fig. 2g) were similarly analyzed and shown to consist of a rod-like structure, as previously described (Fig. 2h)^16^.

The same CLEM strategy (Fig. S2-S5) was applied to precisely locate and visualize the ultrastructure of aSyn-immunopositive inclusions within neurites (Figs. 3, 4, S6f, S7). A 3D TEM tomogram of one such Lewy neurite (LN) in Donor B-PD contained disordered neurofilaments interspersed with vesicular structures reminiscent of mitochondria or remnants thereof (Fig. 3a, blue arrowheads), autophagosomes^14^ (Fig. 3a, b, white asterisk), and sporadically a dense-core vesicle (Fig. 3a, black arrowhead). Importantly, a transition from order to disorder can be seen in the bottom part of Fig. 3b; structures that look like neurofilaments^16^ (Fig. 3b, white arrowheads) appear to become disrupted over the length of the neurite (Movies 16-17). Such disorganization was not observed in brain samples of aged non-neurological control subject; the paths of neurofilaments are far more sharply delineated (Fig. 3c, d), and mitochondria within the neurites exhibit intact cristae (Fig. 3d, black asterisk) (Movies 18-19). Another 3D TEM tomogram of a separate aSyn-immunopositive LN (Fig. S6f) in a different donor (Donor D-PD) was shown to consist entirely of vesicles and membranous structures. A third LN, found in Donor E-PD, contained abundant tubulovesicular structures interspersed with filaments (Fig. 4).

Serial block-face scanning electron microscopy (SBFSEM)^11^ of corresponding tissue from one of the same brain donors (Donor B-PD) as used for CLEM, provided 3D reconstructions of aSyn-immunopositive inclusions across volumes spanning tens of microns (Fig. 5, Movie 20), resulting in a comprehensive 3D view of these aggregates in tissue at a spatial resolution of ∼16 nm. The SBFSEM data revealed cytoplasmic inclusions of aggregated intracellular material (Fig. 5, Movie 20) and the presence of many mitochondria (Fig. 5a-b, orange arrows) and other organelles resembling autophagosomes and lysosomes (Fig. 5a, b; aqua arrows), in agreement with Nixon and colleagues^14^. A representative inclusion identified by SBFSEM was imaged at higher resolution (100-200 nm) by correlative TEM to define their ultrastructure more clearly. The higher-resolution TEM images showed the analyzed inclusions to be comprised of membrane fragments (Fig. 6). These have clearly distinguishable lipid bilayers and are similar in appearance to the membranes of organelles, such as mitochondria and the plasma membranes of cells. One inclusion also exhibited dysmorphic, elongated mitochondria at its immediate periphery, either clustered with other membrane fragments and vesicles (Fig. 6b) or encased in distinct cellular compartments (Fig. 6c) in a state of what appeared to be partial degradation. Other mitochondria were more centrally located in the aggregate and surrounded by membrane fragments (Fig. 6d).

Clusters of membrane fragments, vesicles and structures resembling dysmorphic mitochondria (Fig. 6f), omegasomes (autophagosome precursors; Fig. 6f, g, black arrowheads) and lysosomes (Fig. 6f, g, pink arrowheads), were located towards the center of the inclusion. Filamentous structures were observed, *e.g.*, cytoskeletal filaments were evident in regions within the same tissue section but distal to the LB (Fig. 6i, j). Furthermore, most mitochondria within the inclusion (Fig. 6b, c, d, f) were dysmorphic compared to mitochondria found within the cytoplasm of intact cells of the same tissue section, yet distal to the inclusion (Fig. 6e, i).

To clarify the identities and distributions of vesicular structures found within Lewy pathology, multiple labeling experiments followed by 3D gated STED microscopy was applied (n = 14 PD donors in total). Lysosomal and mitochondrial markers were chosen for STED investigations since lysosomal-type structures and mitochondria were observed in the aSyn-immunopositive inclusions using EM. Morphologies of LB/LN were classified based on the presence or absence of an outer layer of aSyn phosphorylated at Serine 129, here termed p-aSyn. LB with a uniform distribution of p-aSyn reactivity throughout the structure were observed in hippocampus and SN, and showed widespread immunoreactivity for LAMP-1 as well as VDAC1-immunopositive mitochondria (Fig. S13), along with some inclusions showing empty vacuoles that may represent autophagic vacuolar-like structures (membrane-enclosed, “empty” vesicles) (representative one shown in Fig. S13, f, i, j). The presence of empty vacuoles in these inclusions by STED bears similarity to separate aSyn-immunopositive inclusions identified by CLEM (Figs. 4, S12). LB and LN with an outer layer of p-aSyn immunoreactivity were observed predominantly in the SN, and displayed a peripheral clustering of VDAC1-immunopositive mitochondria (Figs. 5d, i, n; S13; Movie 21).

The STED images of LB with uniform distribution of p-aSyn support the CLEM data (Figs. 1, 2, 4, S6, S9, S10) and SBFSEM data (Fig. 5, Movie 20) that demonstrate a relatively even distribution of lysosomal-type structures and mitochondria interspersed. Interestingly, in LB with a p-aSyn positive outer layer, we also observed a shell of mitochondria directly surrounding some of the inclusions (Fig. 5a-b), corroborating our SBFSEM data (Fig. 5, Movie 20), correlative SBFSEM/TEM (Fig. 6) and 2D CLEM data (Fig. S12). Overall, the STED data generated on brain samples from 14 PD donors confirm that different Lewy inclusion morphologies (with and without an outer layer of p-aSyn) contain organelles or organellar remnants, including mitochondria and lysosomes.

To cross-validate the high lipidic content of aSyn-immunopositive inclusions as observed in our TEM tomograms, 10 µm-thick cryostat sections were collected from tissue blocks of the same SN region from the same brain donor (Donor B-PD; Table 1) and the CA2 region of the hippocampus obtained from another late stage PD brain donor (Donor A-PD; Table 1). These cryostat-cut tissue sections were then co-stained for lipids (Nile Red) and aSyn (immunohistochemistry labeling; Methods). In both cases, projections of the image stacks collected by confocal fluorescence microscopy showed co-localized staining of lipids and aSyn in the LB examined (Fig. S14), supporting the concept of a high membrane content therein.

LB were examined by three additional methods to confirm the membranous lipid content shown by confocal fluorescence microscopy (Fig. S14), and thus corroborate the presence of cellular/organellar membranes indicated by the EM data (Figs. 1a-c, 2-6, S6a-c, e-f, S9-S12). The first method combined coherent anti-Stokes Raman scattering (CARS)^17^ with subsequent immunofluorescence staining and confocal laser scanning microscopy (CLSM) in correlative measurements, herein referred as correlative CARS-IF. CARS is a nonlinear optical imaging method that allows the label-free identification of the chemical composition of lipids and proteins in tissue at a resolution of 300 nm. Cryostat sections cut from tissue blocks taken from the same brain regions of the brain donors as used for the EM and LM studies were investigated, namely the CA2 region of Donor A-PD, and the SN region of Donor B-PD. CARS detected a high lipid content throughout aSyn-labeled inclusions: in areas that showed a higher aSyn signal, we observed a higher lipid signal as compared to the surrounding tissue (Fig. S15, a-d) in the correlative tissue sections.

The second method used to confirm the presence of lipids, combined high-definition Fourier transform infrared (FTIR) spectroscopic imaging with subsequent correlative immunofluorescence staining to detect aSyn and confocal laser scanning microscopy (CLSM), herein referred as correlative FTIR-IF. FTIR imaging exploits the fact that the absorption of mid-infrared light waves by chemical bonds (*e.g.,* C=O, C-H, N-H) depends on the chemical environment of the bonds, *i.e.,* the presence and composition of specific biomolecules within cells or tissue (*i.e.*, lipids, protein, DNA)^18^. FTIR therefore does not require the use of labels. The FTIR-IF results confirmed that aSyn-immunoreactive inclusions are rich in lipids (Fig. S15, e-h). The CARS and FTIR lipid and protein profile of neurons in non-neurological controls showed less lipids and proteins than the surrounding tissue whereas LBs showed an increase in lipid and protein profile in the SN and CA2 (Fig. S16). With these measurements, we showed that the increase in the lipid and protein signals in these inclusions were indeed attributable to the actual inclusion itself.

Finally, liquid chromatography-mass spectrometry (LC-MS) and lipidomics analyses were applied to LB isolated by laser-capture microdissection microscopy (LCM). LB identified by aSyn-immunostaining were extracted from 7 µm-thick cryostat-cut tissue sections of SN (∼3050 LB) and CA2 (∼2700 LB) of the same PD brain donors (Donors B-PD and A-PD, respectively; Table 1) using LCM. The subsequent analysis confirmed the high cell membrane related lipid content indicated by other methods, with the mass spectra showing strong peaks corresponding to sphingomyelin (SM) and phosphatidylcholine (PC) for aSyn-immunopositive inclusions isolated from both the SN and hippocampal CA2 sector (Fig. S17b, c). Similar peaks were observed in myelin-rich/lipid-rich regions dissected from the corpus callosum of a non-neurological control brain donor, and cells of the dentate gyrus (DG) from the hippocampus of Donor A-PD (Fig. S17a, d). The SM/PC profile is not specific for LB, which contain abundant membranes that originate from the cell itself; therefore, it is not surprising that their lipid profile would be similar.

Together, these three orthogonal methods confirm that Lewy pathology contain both aSyn and lipids in close proximity and show that they are rich in lipids found in other physiological and lipid-rich structures in the brain, *e.g.*, organellar membranes. They fully corroborate the interpretation of the microscopy data, confirming that Lewy pathology contains aSyn, lipids, lysosomes and mitochondria.

## Discussion

Our results using advanced electron and optical imaging techniques show that Lewy pathology in brain tissue of PD donors generally consist of a crowded medley of mitochondria, lysosomes and membranous vesicles. These findings support a key role for potentially damaged, distorted organelles in the formation of LB and LN, a major process in the pathogenesis of PD. Our criteria for identification of LB and LN was primarily based on the strong staining for LB509 (anti-aSyn), which is routinely used to identify Lewy pathology, and allowed distinguishing from other brain inclusions such as *corpora amylacea*^19^. Given that each tissue section collected for LM and EM was 100-200 nm in thickness, we could overlay the same tissue features very precisely in one section for LM and the adjacent section for EM. Our CLEM approach thereby left no doubt that the spherical aggregates of similarly reported LB diameters (∼4-25 µm)^9, 20^ at the indicated locations of LB found by light microscopy, must indeed correspond to the matching structures in EM. SBFSEM provided 3D reconstructions of aSyn-immunopositive inclusion across volumes spanning tens of microns (Fig. 5, Movie 20) giving a comprehensive 3D view of these inclusions in tissue at a spatial resolution of ∼16 nm. Comparatively smaller sub-regions of interest were investigated at higher resolution (100-200 nm) by correlative SBFSEM/TEM (Fig. 6) and in 3D by TEM tomography (Figs. 1-3, S7) and 2D (Fig. 4, S8-S12) using CLEM (Figs. S2-S4). Together, this led to the discovery of ultrastructural membranous features in Lewy pathology; contrary to expectations, we detected primarily filamentous morphology in only three out of 16 aSyn-immunopositive inclusions (Supplementary Table 1). The CLEM, SBFSEM and TEM was complemented by confocal IF microscopy, 3D reconstruction of multi-labeling STED microscopy data (Figs. 5, S13), correlative CARS-IF and FTIR-IF imaging (Fig. S15-S16). These methods confirmed that lipids and markers for lysosomes and mitochondria are present in aSyn immunoreactive LB and LN, and further provided evidence for crowding of organelles in such.

Our CLEM approach has permitted, for the first time, an unbiased localization at the nanometer scale of Lewy pathology using aSyn immunohistochemistry in tissue sections compatible for both LM and high-quality EM. Given the distinctive morphology of historically-identified LB within the monochrome and crowded cellular landscape by EM, our results help explain why in past studies only distinctly filamentous inclusions could be identified by EM alone. The other aSyn-immunopositive inclusions containing a crowded medley of lipids and organelles are more difficult to distinguish from the cellular background in TEM images alone, and can only be found when using CLEM, as demonstrated in our study.

It is important to note that clarifying the nature of the observed filaments in aSyn-immunopositive inclusions, *i.e.*, distinguishing abundant cytoskeletal filamentous structures such as neurofilaments and other kinds of filaments from aSyn filaments, was not possible in this study. For example, aSyn filaments in brain tissue are reported as similar in diameter to neurofilaments, measuring at 5-10 nm or 9-12 nm while neurofilaments are 10 nm^3, 9, 21, 22^. Immunogold or related staining procedures for immunoelectron microscopy on postmortem brain tissues can provide sufficient resolution to localize filaments using immunogold markers. However, clarifying whether immunogold localizes truly to filaments or to the aggregated (proteinaceous) material surrounding filaments in brain tissue extracts, can be challenging^9, 22^. A further complicating factor is the filament extraction process itself that is based upon protocols optimized for the extraction of paired helical and straight filaments of Alzheimer’s diseased brains^9^. When such protocols were applied to DLB brains with concomitant tau and Aβ pathology^23, 24^, distinguishing Aβ/tau filaments from aSyn filaments has so far not been done, and would be challenging. Even considering the scant literature on aSyn immunoelectron microscopy in PD postmortem brain tissue^3^, the protocols commonly used for tissue processing for EM preclude making reliable conclusions regarding immunopositive aSyn clusters with respect to the presence and identity of filaments vs. actual aSyn filaments within Lewy pathology. This issue is further complicated when considering that mono- and multimeric forms of aSyn might simultaneously be present in this crowded intracellular environment^25^.

Besides being useful to distinguish Lewy pathology from the crowded landscape typically seen in brain tissue electron micrographs, our CLEM approach was crucial to distinguish LB from age-related changes, in particular *corpora amylacea* (CA). CAs and LB are both similar in size, shape and brain tissue localization in aged individuals (Fig. S18). Lewy himself noted the conundrum of distinguishing between the two^26^, and it has remained a challenge throughout the past century. It cannot be excluded that some of the structures previously analyzed by EM and published as a characterization of Lewy structures were in fact CA^19^.

While previous studies also indicated lipid content in LB^24^ *in situ* or as potential membranous contamination in brain extracts considered to contain LB^23^, our nanoscale imaging-based discovery of a crowded medley of membrane fragments, mitochondria, vesicular structures, including some that resemble lysosomes and autophagosomes, intermingled with non-fibrillar aSyn in multiple pathological inclusions, provokes new theories about the mechanisms contributing to the formation of Lewy pathology in PD. A recent study by Grassi and colleagues^27^ has shown that non-fibrillar aSyn species produced from a partial autophagosomal digest appear to disrupt mitochondria in both postmortem brain tissue and cell culture, which corroborates our CLEM data (Figs. 1-3, 5a-b, 6b-d, S6). Although the impact on mitochondrial maintenance for specific genes mutated in idiopathic PD (except for *SNCA*) is difficult to specify^28^, our findings of mitochondria around and within LB, many of which appear distorted and clustered together or in a damaged state, indicate potential mitochondrial instability or dysfunction in certain neurons in PD. Many of the Mendelian genes (*SNCA, LRRK2, VPS35)*, *GBA* (the single risk gene for PD), and various risk *loci* are associated with autophagy and lysosomal degradation^29, 30^. Mitochondrial homeostasis is known to be influenced by aSyn, which interacts with mitochondria-associated ER membranes, and is suggested to play a role in disrupting autophagy, endosomal transport, and ER traffic to the Golgi^31^. Furthermore, the recently described novel pathogenic variant in LRP-10 described in Donor B-PD has been linked to disturbance of intracellular membrane trafficking in PD^32^. Our finding that mitochondria co-localize with membrane fragments and membranous features reminiscent of autophagosomes within LB, supports the hypothesis that dysregulation of intracellular degradation pathways for proteins and entire organelles and disturbed intracellular membrane trafficking is a determining factor in PD. This crowded medley of organelles may also represent a result of the cell’s efforts to sequester problematic lipid-based contents associated with aSyn into an aggresome-like structure. Indeed, McNaught and colleagues have shown that LB are immunoreactive for several markers of aggresomes^33^, deposits that form in response to cytoplasmic accumulation of misfolded proteins^34–36^. The formation of such LB may hence arise from an aggresome-like process^33, 36^. Furthermore, studies have shown that crowding of aSyn on membranes not only changes the morphology of vesicles and mitochondria, but can also catalyze the formation of aSyn filaments^37^, which may explain the presence of distorted mitochondria and vesicles in the LB analyzed in our work.

Our correlative TEM tomograms show clearly that LN are also primarily composed of membrane fragments, dysmorphic mitochondria and structures reminiscent of lysosomes and autophagosomes, as well as cytoskeletal building blocks. To the best of our knowledge, such an observation has not been reported before for LN. The cytoskeletal abnormalities recorded by CLEM tomography in two regions of a LN and two regions of another LN of a different brain donor using 2D CLEM support the idea that neurofilaments become disrupted, possibly through proximally experiencing an increase of oxidative stress^38, 39^. The 3D STED data also showed that LN contain many mitochondria and lysosomes. One could speculate that a crowded medley of damaged organelles and proteins would be sufficient to disrupt axonal trafficking.

The precise form and location of aSyn within the LB and LN here observed by CLEM is not known. Since all Lewy features studied here were identified by their high content of aSyn, but many did not contain any filamentous material, aSyn must be present in these LB and LN in a different form. One possible interpretation is that aSyn may be acting as membrane tether^40^, bridging different mitochondrial membrane patches and leading to excessive adhesion between mitochondrial membranes, which could have led to mitochondrial damage, membrane disruption and formation of Lewy pathology – consisting of fragmented organellar membranes, each excessively decorated with aSyn. Alternatively, aSyn might have disrupted the membrane integrity, leading to fragmented organelles, which subsequently clustered into LB.

Either way, our data indicate that aSyn may modulate the compartmentalization and function of membranes and organelles in LB-affected cells, prompting a new hypothesis about the role of aSyn in the formation of Lewy pathology in PD (Fig. 7).

**Figure 7.**
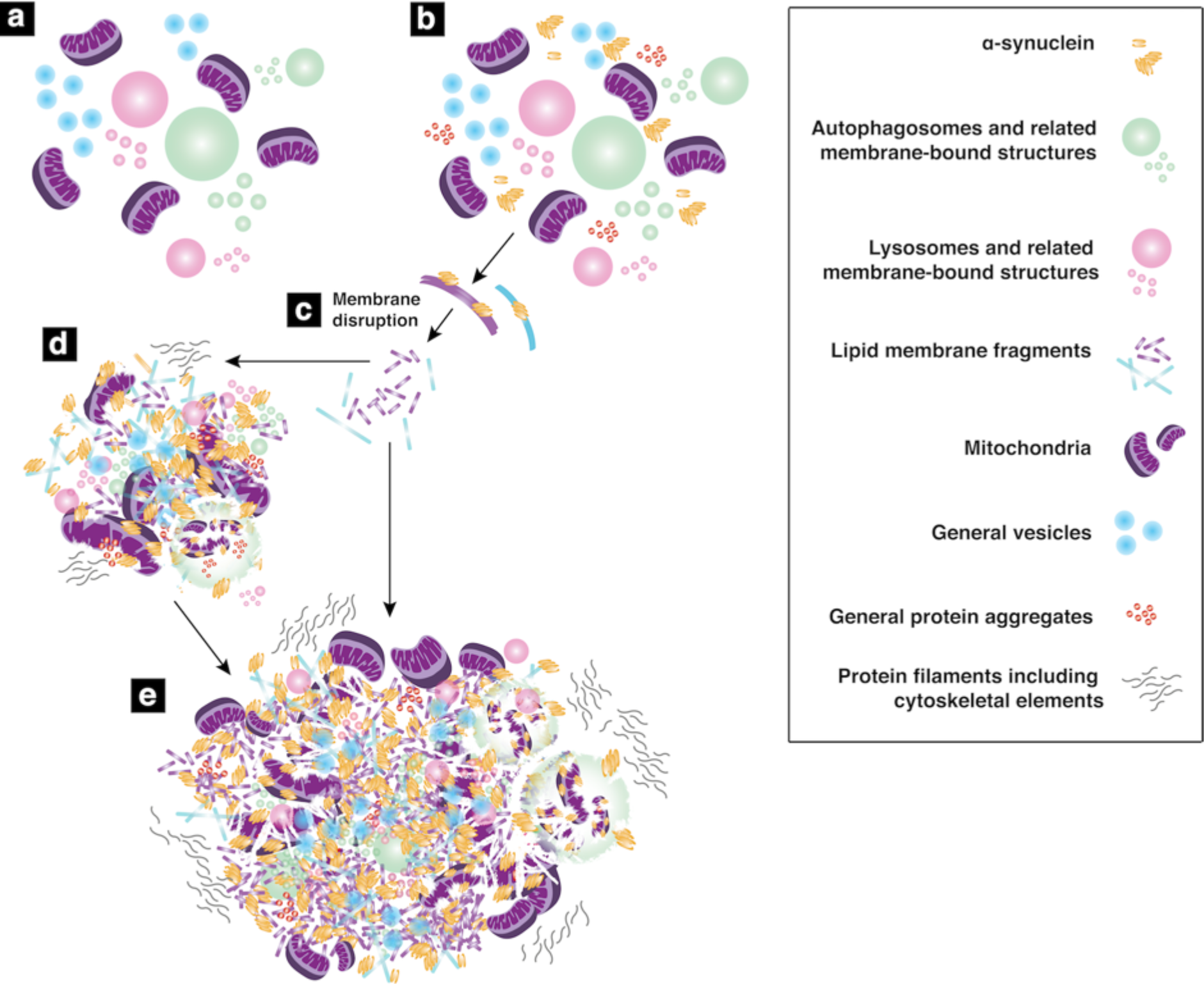
Hypothetical mechanism for the formation of membranous Lewy pathology in PD. **(a)** Organelles including mitochondria as they exist physiologically in the cell, and **(b)** in the presence of pathologically aggregated or modified aSyn (e.g., phosphorylated at Ser129, oxidated, truncated, etc.) together with other protein aggregates. Over time, this may lead to **(c)** disruption of organellar membranes and **(d)** further aggregation of organelles and disruption and fragmentation of their lipid membranes. **(e)** Larger clumps of lipid membrane fragments, aggregated proteins, vesicles and other general membrane bound structures, which compact over time in the restricted cellular environment, would give rise to the ultrastructure of the majority of Lewy pathology (membrane-rich) as observed by the CLEM, SBFSEM and STED methods used in this study.

The single LB with radiating filaments identified in our work was interestingly found amongst neuromelanin-containing organelles, which are highly concentrated vesicles, specifically autolysosomes^41^. By contrast, the membrane-rich, aSyn-immunopositive inclusions were found both outside and within neuromelanin-containing organelles.

The physiological role of aSyn in the presynaptic terminal includes remodeling of membrane, clustering of synaptic vesicles and maintaining synaptic vesicle pools, promoting SNARE-complex assembly, and modulating the release cycle of synaptic vesicles^42^. The formation of structures called ‘nanodiscs’ by aSyn and lipids *in vitro*^43^ has been reported. Our findings of membrane-rich Lewy pathology in human brain are similar to observations by EM in a transgenic mouse model that overexpresses human aSyn^44^. Boassa and colleagues had found alterations and enlargements of the presynaptic endomembrane systems in these mice, with presynaptic terminals filled with membrane-bound organelles, including tubulovesicular structures. They proposed that aSyn overexpression is associated with hypertrophy of membrane systems. Our findings of abundant membranes in actual human Lewy pathology supports their animal model.

Another research study in rats demonstrated that aSyn interacts with tubulovesicular/vesicular structures through its amino-terminal repeat region^45^. It is tempting to suggest that these ‘nanodiscs’ and tubulovesicular structures are a rudimentary form of the membrane ‘discs’ observed in Lewy pathology, such as shown in our 3D CLEM tomogram and abundantly visible in our 2D CLEM images of other brain donors. Furthermore, the observation of tailed membrane stacks exhibiting a bilayer separation of ∼6 nm (Movie 12), different than expected for myelin, may point to another role of aSyn in modulating membrane interactions. Intriguingly, certain N-terminal mutations that cause reduced solubility of aSyn *in vitro* lead to the formation of round inclusion bodies in transfected primary rat neurons^46^, but whether membranes or organelles play a role in this model remains to be demonstrated.

In future experiments, it will be interesting to explore how aSyn influences the formation of Lewy pathology in concert with vesicles and cytoplasmic organelles. In light of the presented findings, it is also relevant to explore whether the animal and cellular models of intracellular aSyn accumulation reveal similarities to our observation in human brain, thereby leading to more representative translational models of PD and related synucleinopathies.

Our work provides a new dimension of understanding of Lewy pathology in the human brain, with nanoscale imaging that is cross-validated by orthogonal biophysical methods. We present here a new theoretical model in which lipid membrane fragments and distorted organelles together with a non-fibrillar form of aSyn are the main building blocks for the formation of Lewy pathology (Fig. 7). Our model for the formation of these highly membranous aSyn-immunopositive inclusions is supported by evidence from existing studies that demonstrate aSyn as capable of disrupting mitochondrial membranes, manipulating and organizing membrane components, and inducing membrane curvature to vesicles under certain conditions^27, 43, 47–51^.

Together, our results support the hypothesis of impaired organellar trafficking as a potential driver of pathogenesis in Parkinson’s disease. Further studies are now required to study the morphology of Lewy pathology across different Braak stages of PD, and to analyze the brain tissue ultrastructure in related synucleinopathies. Our findings emphasize the need to consider population heterogeneity of Lewy pathology and that lipids are a major component of them. This has strong implications for the understanding of the pathogenesis of PD. Importantly, it could lead to new approaches for cellular and in vivo models, support the rationale for the development of urgently needed biomarkers such as PET tracers for Lewy pathology, and point to novel drug targets for Parkinson’s and related diseases.

## Methods

### Human Postmortem Brain Tissue Samples

Post-mortem tissue samples from five donors (Donors A - E) with clinical diagnosis PD with dementia (PDD) and one non-neurological control (Donor F-Control), all with ∼5 hours post-mortem delay, were obtained from the Netherlands Brain Bank (NBB, www.brainbank.nl; Table 1) and the Normal Aging Brain Collection (Dept. Anatomy and Neurosciences, VUmc), respectively. Tissues were collected using a rapid autopsy protocol (NBB). All donors had given written informed consent for a brain autopsy and the use of the material and clinical information for research purposes. Detailed neuropathological and clinical information was made available, in compliance with local ethical and legal guidelines, and all protocols were approved by the local institutional review board.

At autopsy, four 0.5cm-thick adjacent brain slices of the mesencephalon and hippocampus (mid) were collected. Cubes of ∼1-2 mm^3^ of the ventral part of the *substantia nigra pars compacta* (SNpc) and hippocampal CA2 regions were dissected and fixed for 6 hours in a mixture of 2% paraformaldehyde/2.5% glutaraldehyde in 0.15 M cacodylate buffer with 2 mM calcium chloride, pH 7.4 and then washed with PBS. One slice of mesencephalon and hippocampus was directly snap-frozen for processing for LCM and subsequent lipidomics and CARS analysis.

The PD brain donors fulfilled the United Kingdom Parkinson’s Disease Society Brain Bank (UK-PDSBB) clinical diagnostic criteria for PD^52^. Neuropathological evaluation was performed on 7 µm formalin-fixed paraffin-embedded sections collected from multiple brain regions according to the guidelines of BrainNet Europe. As is routine for such brain donors, staging of Alzheimer’s disease was evaluated according to the Braak criteria for NFTs^53^, CERAD criteria adjusted for age and Thal criteria^54^. The presence and topographical distribution of aSyn (monoclonal mouse anti-human-α-synuclein, clone KM51, Monosan; Fig. S1) was rated according to Braak’s staging scheme for aSyn^7^ and a modified version of McKeith’s staging system for aSyn (*i.e.*, brainstem, limbic system, amygdala-predominant or neocortical^55^.

### Whole Exome Sequencing and PD Gene Analysis

Postmortem brain tissues from donors used for these EM studies (Table 1) were analyzed by whole exome sequencing (WES) with a focus on the analysis of the PARK genes and additionally some genetic risk factors for dementia. No listed known causative genetic variants were detected in the donors. In Donor B-PD, a variant in the LRP-10 gene, a potential new gene causal for PD that needs yet to be confirmed by others, as previously described^32^. Variants were to be reported if they fulfilled the following four criteria: (1) Variant(s) located within the following list of genes associated with PD or Parkinsonian syndromes: ATP13A2; ATP6AP2; CHCHD2; DNAJC13; DNAJC6; EIF4G1; FBXO7; GBA; LRRK2; PARK2; PARK7; PINK1; PLA2G6; RAB39B; SNCA; SYNJ1; TARDBP; VPS35; VPS13C; MAPT; GRN; TMEM230; POLG; DCTN1; PTRHD1. (2) Possible splicing variants → intronic or exonic variants (synonymous variants have been included) located within 10 bp at the exon-intron boundaries. (3) Exonic variants that have a coding effect (synonymous variants have been excluded). (4) Novel variants or variants present with a minor allele frequency (MAF; below 1%) in the following publicly available databases: NHLBI Grand Opportunity Exome Sequencing Project (ESP) (http://evs.gs.washington.edu/EVS/); Exome Aggregation Consortium Browser (ExAC) (http://exac.broadinstitute.org/); 1000 Genomes (http://browser.1000genomes.org/index.html); dbSNPs (https://www.ncbi.nlm.nih.gov/projects/SNP/); Genome of the Netherlands (GoNL) (http://www.nlgenome.nl/).

### Correlative Light and Electron Microscopy

The CLEM workflow is summarized in Fig. S4, and pictorially shown in Figs. S2-S3. Fixed postmortem human brain tissue (2% filtered paraformaldehyde/2.5% glutaraldehyde in 0.15 M cacodylate buffer with 2 mM calcium chloride, pH 7.4) was washed in cacodylate buffer and kept at 4 °C for 1-2 days. Tissue sections were then collected at 40-60 µm on a vibratome and washed in cold cacodylate buffer, post-fixed in potassium ferrocyanide in cacodylate buffer with osmium tetroxide, washed with double-distilled water, and immersed in filtered thiocarbohydrazide solution. After this second wash step, sections were post-fixed in osmium tetroxide, rinsed again and stained with uranyl acetate at 4 °C overnight. The following day, sections were rinsed, stained with lead aspartate solution at 60 °C, dehydrated in a graded alcohol series on ice, and embedded in Durcupan resin. After resin hardening, small pieces of the resin-embedded tissue (∼ 1 mm x 1 mm) were cut and mounted on standard aluminum pins, then physically cut using a razor blade into a trapezoid shape, which is optimal for the collection of serial tissue sections.

All tissues sections were generated using a physical ultramicrotome (Ultracut EM UC7; Leica Microsystems, Germany) and cut at a thickness of 100-200 nm. They were alternately collected on Superfrost™ Plus glass slides (Thermo Fisher Scientific, USA) for later light microscopy (LM) and EM grids (EMS Diasum, PA, USA) with a carbon-stabilized formvar film for TEM imaging. Slides were processed for immunohistochemistry using mouse anti-aSyn (Invitrogen 180215, LB509 concentrate) diluted 1:500. The sections were etched in a saturated ethanolic potassium hydroxide solution for 5 minutes followed by washing in PBS. Endogenous peroxides were quenched with 1 % hydrogen peroxide in 10 % methanol for 10 minutes followed by blocking in antibody diluent (Dako /S 202230) for 10 minutes. The sections were incubated in primary antibody for 1 hour at 37 °C followed by washing with 0.25 % Triton X-100 in PBS and incubation with the Immpress Reagent Anti-Mouse Ig (Vector/VC-MP-7401) secondary antibody for half and hour at room temperature. Bound antibody complexes were detected using the Permanent HRP Green Kit (Zytomed Systems) with incubation for 3 minutes at room temperature, before counterstaining with hematoxylin, dehydration and coverslipping.

The slides were screened by LM and compared side-by-side to identify Lewy pathology. LM images of selected slides displaying aSyn-immunopositive inclusions were collected at 40x magnification using a Zeiss Axiophot (Carl Zeiss Microscopy) with monochromatic light, or at 60x magnification using a Nikon Ti-E widefield and the images were manually stitched together using Adobe Photoshop CS6 (Adobe Systems Incorporated) or FIJI to create a montage revealing the trapezoid shape of an individual tissue section. The full montage representing the individual tissue section was then cropped to the limits of the tissue borders using an edge-detection lasso tool, which followed the trapezoid shape of the tissue sections. TEM images of EM grids that contained tissue sections immediately adjacent to those on the selected LM slides, were collected either on a Talos 200 keV TEM (FEI, Thermo Fisher, USA) and manually stitched together using Adobe Photoshop, or on a Titan Krios 300 keV TEM (FEI, Thermo Fisher, USA) using the SerialEM^56^ montage option. The resulting EM montage was overlaid with the corresponding LM montage obtained for the alternating tissue sections collected on a glass slide, to define the specific location of the aSyn-immunopositive inclusions in the TEM images (Figs. S2-S5) and guide the collection of subsequent higher resolution images and electron tomography. The collection of serial tissue sections on a single EM grid meant that the same aSyn-immunopositive inclusion was present multiple times and that obstruction of the inclusion by a grid bar was not an issue, since another section of the same grid, where it was not obscured, could be used.

For TEM tomography, samples were imaged at cryogenic temperatures using a Titan Krios (FEI, Thermo Fisher Scientific, USA) equipped with a Quantum-LS energy filter (20 eV slit width) and a K2 Summit direct electron detector (Gatan, Pleasanton, CA, USA) and operated at 300 kV acceleration voltage, or at room temperature on a Talos (FEI, Thermo Fisher Scientific, USA) operated at 200 kV. Tilt series were recorded using the SerialEM software^56^ with a unidirectional tilt-scheme at 2-3 degree increments or a “dose-symmetric Hagen tilt-scheme”. The latter procedure begins at low tilt and then alternates between increasingly positive and negative tilts to maximize the amount of high-resolution information maintained in the tomogram for subsequent subtomogram averaging and 3D color segmentation, and also yields improved tracking between sequential tilt angles^57^. Images for the tilt series were collected at 3° increments over a range between −60° to 60° at a nominal defocus within 6-10 µm.

Tilt series alignment by cross-correlation and patch-tracking followed by 3D reconstruction of unbinned tomograms were performed using *etomo* of the IMOD software^58^. Resulting tomograms were reduced by a factor of 2 in all dimensions. Semi-automatic 3D color segmentation of the tomograms was performed by user-interactive thresholding and volume rendering using the Amira 6.0 software (FEI). Movies of 3D color segmented tomograms were created using Amira 6.0. Movies of reconstructed, non-color segmented tomograms were created using IMOD software. Subtomographic texture analysis was performed using the *Dynamo* software^59, 60^.

### Serial Block-Face Scanning Electron Microscopy and Correlative Transmission Electron Microscopy

Fixed postmortem human brain tissue (2% filtered paraformaldehyde / 2.5% glutaraldehyde in 0.15 M cacodylate buffer with 2 mM calcium chloride, pH 7.4) was washed in cacodylate buffer and kept at 4 °C for 1-2 days. Tissue sections were then collected at 40-60 µm on a vibratome, washed in cold cacodylate buffer, post-fixed in potassium ferrocyanide in cacodylate buffer with osmium tetroxide, washed with double-distilled water, and immersed in filtered thiocarbohydrazide solution. After this step, sections were post-fixed in osmium tetroxide, rinsed again and stained with uranyl acetate at 4 °C overnight. The following day, sections were rinsed, stained with lead aspartate solution at 60 °C, dehydrated in a graded alcohol series on ice, and embedded in Durcupan resin. After resin hardening, small pieces of the resin-embedded tissue (∼ 1 mm x 1 mm) were cut and mounted on standard aluminum pins. The samples on the pins were sputter-coated with gold and platinum in a vacuum system to enhance sample conductivity for SEM imaging, and then directly transferred to the SEM chamber for imaging.

Data were collected using a scanning electron microscope (FEI Quanta200FEG, Thermo Fisher, USA) equipped with a physical microtome (3View, Gatan, Pleasanton, CA, USA) inside the microscope observation chamber. An accelerating voltage of 3.5 keV, a spot size of 3, a scanning speed of 2 µsec/pixel and the high vacuum mode were used. After each iterative removal of an ultrathin slice (70 nm thick) by a diamond knife within the SEM chamber, the surface of the remaining block (specimen) was imaged. Images with 8192×8192 or 4096×4096 pixels were collected at 7-10 nm/pixel along both the x- and y-axes using the Digital Micrograph software (Gatan). Image series of regions of interest were further processed, digitally aligned and reconstructed into 3D z-stacks/tomograms using the TrakEM2 module of Fiji (https://fiji.sc).

To enable correlative TEM, once a region of interest containing inclusion bodies was identified and partly imaged by SEM, the sample was removed from the SEM chamber and cut using a physical ultramicrotome (Ultracut EM UC7; Leica Microsystems, Germany). The resulting 30-50nm-thick slices obtained were sequentially collected on EM grids (EMS Diasum) with a carbon-stabilized formvar film, and imaged at room temperature (RT) using a Philips CM10 electron microscope operated at 80 kV. Electron micrographs were recorded on a 2,048×2,048-pixel charge-coupled device camera (Veleta; EMSIS GmbH, Germany). Color annotation of the resulting 2D micrographs was performed manually using Adobe Photoshop CS6 (Adobe Systems Incorporated).

### Lipid-aSyn Co-Staining and Fluorescence Imaging

Tissue sections (10 µm thick) of a snap-frozen tissue slice of the hippocampus (including CA2) of Donor A-PD and the SN of Donor B-PD were cut using a cryostat (Leica) collected on glass slides at −18 °C, shipped on dry ice from the VUmc to C-CINA, and subsequently stored at −80 °C. Immediately after removal from −80 °C, slides were fixed with 4% paraformaldehyde (PFA) for 30 minutes in a humidity chamber. They were then rinsed in PBS and treated with 0.5% Triton X-100 in PBS, washed with PBS and incubated for 2 hrs at RT with a primary antibody to aSyn, LB509 (amino acid 115-122; abcam ab27766). After washing with PBS, slides were incubated for 1.5 hrs at RT with a cocktail of secondary fluorescence-conjugated antibody Alexa488 and DAPI (4’,6-diaidino-2-phenylindole, dilactate; BioLegend) to visualize nuclei. Slides were washed with PBS, and then treated with Nile Red stain (Sigma 19123) for 10 min in the dark. Nile Red powder was originally diluted to 0.5 mg/ml in acetone to make a stock solution, and a fresh working solution was prepared every time by diluting an aliquot of this stock 1:200-fold in 75% glycerol. Stained slides were washed in 75% glycerol, treated with Sudan Black in the dark, rinsed in PBS and mounted in Mowiol coverslip mounting solution. They were allowed to dry in the dark overnight, and then stored at 4°C in the dark prior to imaging.

Confocal fluorescence images (1024 × 1024 pixels) were mainly acquired at a magnification of 40x, using a point-scanning confocal microscope (CLSM Leica TCS SPE with a DMI4000 microscope) equipped with advanced correction system (ACS) objectives and the solid-state laser system: 405 nm (DAPI), 488 nm (Alexa488), and 635 nm (Nile Red). Composite images and co-localizations were calculated and created using standard tools in the Imaris software (Bitplane AG) and final figures were composed in Adobe Photoshop CS6 (Adobe Systems Incorporated). Untreated tissue samples were checked by LM before and after staining/labeling to look for auto-fluorescence, as this could interfere with label detection, and then by confocal laser scanning microscopy (CLSM) to visualize greater detail in 2D and 3D. As a further control, tissue sections were treated with the secondary fluorescence antibody alone, without the primary antibody, and examined by LM and CLSM to check for unspecific labeling.

### Co-labeling for STED Microscopy

Multiple labeling experiments were performed on formalin-fixed, paraffin-embedded 20µm-thick midbrain and hippocampus sections, using markers for organelles and aSyn. For heat-induced antigen retrieval, sections were placed in sodium citrate buffer (pH 6.0) in a steamer at 90-99 °C for 30 min. Antibodies against VDAC-1/Porin (Abcam ab14734), LAMP-1 (Abcam ab24170) and S129-phosphorylated α-synuclein (pSer129 Ab 11A5) directly labeled with AlexaFluor 488 were used. Abberior STAR 580 and STAR 635P fluorophores (Abberior, Bioconnect) were used as secondary antibodies. Nuclei were visualized by DAPI staining (Sigma).

STED microscopy was performed on a Leica TCS SP8 STED 3X microscope (Leica Microsystems). Sections were irradiated with a pulsed white light laser at wavelengths of 499, 587, and 633 nm. A pulsed STED laser line at a wavelength of 775 nm was used to deplete the Abberior fluorophores (580, 635P), and a continuous wave (CW) STED laser with a wavelength of 592 nm was used to deplete the Alexa 488 fluorophore. Further, to obtain confocal images of the DAPI signal, sections were irradiated with a solid-state laser at a wavelength of 405 nm. The DAPI signal was not depleted. All signals were detected using a gated hybrid detector in counting mode. Images were acquired using a HC PL APO CS2 100 × 1.4 NA oil objective lens, and the resolution was set to a pixel size of 20 nm x 20 nm. Finally, deconvolution was performed with Huygens Professional (Scientific Volume Imaging; Huygens, The Netherlands). Images were adjusted for brightness/contrast in ImageJ (National Institute of Health, USA), and final figures were composed using Adobe Photoshop CS6 (Adobe Systems Incorporated).

### Liquid Chromatography Mass Spectrometry and Lipidomics of Microdissected Tissues

Cryostat-cut tissue sections (7 µm) of the hippocampus (mid) of Donor A-PD, the mesencephalon of Donor B-PD, and the corpus callosum of a non-neurological control donor (Donor F-Control) were collected at −18 °C. Sections were stained with haematoxylin (Sigma) for 1 min, washed under tap water for 5 min, quickly washed in sterile water, then stained with Eosin for 10 sec. They were washed in 96% EtOH for 30 seconds, 100% EtOH for 30 sec, then air-dried under a chemical fume hood prior to laser-capture microdissection (LCM). For the second LCM run, adjacent 7µm-thick sections of the Donor A-PD hippocampus were immunostained with an antibody against aSyn FL-140 (sc-10717, Santa Cruz; dilution 1:2000) for 30 min after fixation with 96% EtOH. After rinsing with PBS, the immunostaining was visualized with Envision detection systems peroxidase/DAB (DAKO). Sections were rinsed again with Tris-HCl (pH 7.4) and running tab water, subsequently air-dried and stored at 4 °C.

LCM was used to obtain approximately 3000 aSyn-immunopositive inclusions from the CA2 of Donor A-PD and 2700 Lewy bodies from the SN of Donor B-PD. In both cases, these inclusions were laser-cut from inverted adjacent tissue polyethylene naphthalate (PEN) membrane slides positioned on the stage of a LMD6500 (Leica Microsystems) microscope, collected in the cap of either 0.2 or 0.5 ml Eppendorf tubes, and kept on ice until further processing for mass spectrometry. Dentate gyrus (DG) region was also laser-cut from hippocampus of Donor A-PD as a additional control, and myelin-rich regions were laser-cut from the corpus callosum (myelin-rich, hence lipid-rich) of a non-neurological control brain donor.

For mass spectrometry, 40 – 60 µl of chloroform:methanol 2:1 (v/v) was carefully added to the inverted caps and shaken gently to dissolve the collected aSyn-immunopositive patches. The closed tubes were inspected using a magnification glass to ensure that there was no undissolved material. When this was the case, they were analyzed by liquid chromatography mass spectrometry (LC-MS). A Dionex Ultimate 3000 RSLC nanoUPLC system with Reprospher 100 Si column (3 µm, 150 × 0.4 mm, 100 Å, Dr. Maisch GmbH, Ammerbuch-Entringen, Germany) was used to separate the isolated material by normal phase LC. The column temperature was set to 40 °C with a flow rate of 10 µL/min. Mobile phase A was isopropanol:hexane:100 mM ammonium carboxylate 58:40:2 (v/v/v) and mobile phase B was isopropanol:hexane:100 mM ammonium carboxylate 50:40:10 (v/v/v)^61^. After an initial phase at 40% B for 5 min, the gradient was ramped up from 40% to 100% B over 25 minutes and followed by a steady phase at 100% B for 5 min.

To identify the eluting peaks, the capillary nanoUPLC was connected to a Waters Synapt G2 HRMS mass spectrometer. The mass signals and their fragments were obtained by a MS scan and a survey MS/MS scan method. A standard off/axis ESI source was used as an atmosphere-vacuum interface. The spray voltage was set to 2.8 kV, the desolvation temperature was set to 200 °C, and the source temperature was set to 100 °C. The MS/MS spectra were obtained using mass-dependent collision energies. Waters Masslynx and Progeenesis QI software was used to evaluate the data.

### Correlative CARS/FTIR and Immunofluorescence Imaging

Cryostat-cut tissue sections (10 µm) of the CA2 of Donor A-PD, SN of Donor B-PD and white matter of Donor F-Control were collected at −18 °C in the same manner as prepared for confocal immunofluorescence (IF) imaging, and dried under a stream of dry air at RT before coherent anti-Stokes Raman scattering (CARS) or Fourier transform infrared spectroscopy (FTIR) imaging. No stain was applied prior to imaging. Only data from Donor A-PD is shown (Figs. S15-S16).

CARS images were acquired using a commercial setup consisting of a picosecond-pulsed laser system that generates two synchronized beams collinearly aligned in an inverted confocal microscope (TCS SP5 II CARS; Leica Microsystems, Heidelberg, Germany). In this setup, a fraction of the fundamental light of an Nd:YVO4 (HighQ Laser, Rankweil, Austria) at 1064 nm is coupled into the microscope and used as a Stokes beam in the CARS process. The frequency-doubled output (532 nm) is used to synchronously pump an optical parametric oscillator (picoEmerald; APE, Berlin, Germany), tunable in the 780–960 nm range. The laser beams are focused into the tissue by an HCX IRAPO L water immersion objective (25x/0.95 W CORR; Leica Microsystems). The forward-detected CARS signal is measured via a non-descanned detector. The mean laser power was measured at the tissue position and found to be 28 and 21 mW at 816 and 1064 nm, respectively. A typical pixel dwell time of 32 µs per scan was selected (31 s per image, 1024 × 1024 pixels covering up to 300 × 300 µm sample area, pixel resolution 300 nm^62^. CARS images of tissues were measured at 816 and 806 nm, which correspond to 2850 cm^-1^ (lipids, CH_2_) and 2930 cm^-1^ (proteins, CH_3_). The lipid and protein distribution profiles in aSyn-immunopositive inclusions were calculated using the Image Processing and Statistic toolboxes of Matlab (The Mathworks, Inc., Mass., USA).

For the FTIR measurements, infrared hyperspectral data acquisition was performed in transflection mode using an Agilent Cary 620 microscope with an Agilent Cary 670 spectrometer (Agilent, California, USA). Spectral data were collected by an MCT focal plane array detector with 128 × 128 elements, providing a field of view (FOV) of approximately 422 µm x 422 µm with a 25x magnification. The spectral data were collected from 3700 - 950 cm^-1^ with a spectral resolution of 4 cm^-1^. Fourier transformation was performed with a Mertz phase correction, Blackman-Harris 4-term apodization, and a zero filling of 2. A high numeric aperture of 0.82 was used^63^. Tissue sections were prepared on MirrIR low-e- slides (Kevley, Chesterland, USA) for the transflection (reflection-absorption) measurements. The second derivative that minimizes the effects of the standing wave artifact was tested in addition. This resulted in the same spectral band positions. Therefore, the vector-normalized spectra were used.

The resulting raw spectral maps were pre-processed using the previously described workflow^64^. Strong artifacts possibly arising from cracks or folds in the tissue were eliminated by quality control based on the signal-to-noise ratio and the integral of the amide I band. The remaining spectra were subjected to a Mie and resonance-Mie correction based on Extended Multiplicative Signal Correction, EMSC^65^ in the wavenumber range from 3100 to 950 cm^-1^. The correction was performed with 30 iteration steps. Higher numbers of iteration steps (up to 100) were tested but the resulting spectra did not show further variances.

For immunofluorescence staining, after CARS or FTIR imaging tissue sections were kept on the slides and fixed with 4% formaldehyde for 30 min. Slides were rinsed and shipped in PBS, then treated with 0.5% Triton X-100 in PBS for 10 min. Slides were rinsed with PBS before incubation for 2 hrs at RT with an antibody targeting phosphorylated aSyn, phospho-S129 (pS129; abcam ab59264). After washing with PBS, sections were incubated for 1.5 hrs at RT with a secondary fluorescent antibody (Alexa488), rinsed again with PBS and applied with the next primary antibody to aSyn, LB509 for another 2 hrs at RT. After rinsing with PBS, sections were incubated for 1.5 hrs at RT with a secondary fluorescent antibody (Alexa647), rinsed in PBS, and applied with Sudan Black for 30 min. After rinsing again in PBS, they were finally mounted in Mowiol and allowed to dry in the dark overnight, subsequently being stored at 4 °C.

Brightfield and immunofluorescence images corresponding to each CA2 region imaged by CARS or FTIR were collected at 10x magnification using a confocal microscope (Leica TCS SP5 CARS with a DMI3000). Images from brightfield/immunofluorescence and CARS or FTIR were overlaid and aligned to each other based on the morphology of tissue edges and lipofuscin deposits that appeared as black granules in the cell soma. Images were collected using an Ar-laser with 488 nm (Alexa488) excitation. An overlay of CARS or FTIR images with their counterparts showing aSyn immunofluorescence was performed using Adobe Photoshop CS6 (Adobe Systems Incorporated).

## Acknowledgements

We are grateful to the individuals who participated in the brain donation program and their families, making this study possible. We thank Sabine Ipsen from the University Hospital Basel and Sandrine Bichet from the Friedrich Miescher Institute for training and assisting with the immunohistochemistry, Ariane Fecteau-LeFebvre for EM maintenance, Allert Jonker for help with preparing cryostat-cut tissue sections, Prothena (South San Francisco, CA, USA) for providing the pSer129 11A5 antibody, Advanced Optical Microscopy Core O|2 (www.ao2m.amsterdam) for support with STED imaging, Daniel Mona for help with labeling antibodies, Paul Baumgartner and Karen Bergmann for administrative help, and Shirley Müller for carefully proof-reading and editing the manuscript. S.H.S. was supported by the Roche Postdoctoral Fellowship (RPF) program; this work was in part supported by the Swiss National Science Foundation (SNF Grant CRSII3_154461) and the Synapsis Foundation Switzerland.

## Author Contributions

S.H.S. performed CLEM/TEM and tomography, SBFSEM imaging and 2D/3D color segmentations, analyzed data, and wrote the manuscript; A.J.L. performed CLEM/TEM and tomography; C.G. and A.G.M. trained and supported S.H.S. and A.J.L with SBFSEM and CLEM tissue preparation and imaging; Jü.H. screened LM slides and analyzed LM data of CLEM for localizing Lewy pathology; Jü.H., G.S., and A.J.L. designed and optimized the staining for CLEM and localization of Lewy pathology in LM data with S.H.S; W.vdB., T.M. and E.H. performed STED imaging; P.P.N. trained and supported A.J.L. in sample preparation, data collection, and image processing for TEM tomography; K.N.G. and J.W. assisted with TEM tomography; R.S. and S.H.S. optimized and performed lipid and aSyn co-staining and confocal imaging; D.C. performed sub-tomogram analysis; A.I. processed tissue samples collected at the autopsy room, prepared cryostat tissue and sectioned paraffin embedded tissue; Y.dG. prepared cryostat tissue for CARS and FTIR imaging; A.J.M.R., and W.vdB. performed rapid autopsies of PD brain donors and controls, collected brain tissue and preformed neuropathological assessment; W.vdB. performed neuroanatomical dissections, and performed laser-capture micro dissection with S.H.S.; A.D.P. performed Raman imaging tests; J.E., A.S. and Joerg H. performed LC-MS analysis; D.N., S.F.EM. and K.G. performed and analyzed CARS imaging; F.G. performed FTIR imaging; M.Q., W.F.J.vIJ. and V.B. provided whole exome sequencing and genetic analysis of PD brain donors; B.B. provided technical input to EM analysis of brain tissue in neurodegeneration; S.F. provided expertise in neuropathology, differentiation of Lewy pathology *vs.* CA, and provided optical microscopy data of CA and Lewy pathology in same tissues; M.B., H.S., W.vdB., and M.E.L. designed research, analyzed and interpreted the data, and contributed to writing the manuscript.

## Supplementary Information

**Supplementary Table 1.** Lewy pathology identified and characterized using CLEM show heterogeneity of inclusion types with majority containing mostly an organellar medley.

| Tissue Localization | Membrane fragments and organelles<sup>a</sup> | Undefined filaments<sup>b</sup> | Undefined proteinaceous inclusions<sup>c</sup> | Undefined proteinaceous core<sup>d</sup> | Data shown in | Donor |
| --- | --- | --- | --- | --- | --- | --- |
| Hippocampal (CA2) | +++ | * | 0 | 0 | Fig. 1a, 2a, 2c | A |
| Hippocampal (CA2) | +++ | 0 | 0 | 0 | Fig. S6a, 2b, 2d | A |
| *Substantia nigra* | +++ | * | 0 | 0 | Fig. 1b | B |
| *Substantia nigra* | +++ | * | ++ | 0 | Fig. S6b | B |
| *Substantia nigra* | +++ | * | 0 | 0 | Fig. S6c | B |
| *Substantia nigra* | ++ | ** | 0 | n/a | Fig. 3, S7 | B |
| *Substantia nigra* | ++ | *** | + | X | Fig. 1d, S8 | C |
| *Substantia nigra* | +++ | 0 | + | 0 | Fig. 1c | D |
| *Substantia nigra* | ++ | ** | 0 | 0 | Fig. S6e | D |
| *Substantia nigra* | + | ***⊙ | ***⊙ | 0 | Fig. S6d | D |
| *Substantia nigra* | +++ | ** | 0 | 0 | Fig. S9 | D |
| *Substantia nigra* | +++ | ** | 0 | 0 | Fig. S10 | E |
| *Substantia nigra* | +++ | **⊙ | **⊙ | 0 | Fig. S11 | D |
| *Substantia nigra* | +++ | ** | 0 | 0 | Fig. S12 | E |
| *Substantia nigra*<sup>N</sup> | +++ | 0 | 0 | n/a | Fig. S6f | D |
| *Substantia nigra*<sup>N</sup> | +++ | ** | 0 | n/a | Fig. 4 | E |
| *Substantia nigra*<sup>N</sup> | ++ | ** | +++ | n/a | Fig. 3a,b | B |

<sup>a</sup> “+, ++, +++” indicate increasing level of abundance
<sup>b</sup> Not possible to distinguish between aSyn filaments and neurofilaments, or other filaments of similar diameter and appearance. \*, \*\*, \*\*\* indicate increasing level of abundance of such filaments, whereas “0” indicates absence. Indicated by pink arrowheads (Figs. 1, S5).
<sup>c</sup> Unclear what proteins the proteinaceous inclusions contain or what physical form they have. They appear as cloud-like clumpy dark patches by EM (blue arrowheads, Figs. 1, S5).
<sup>d</sup> Reminiscent of the previously published EM micrographs of Lewy pathology. 0, absent; X, present; n/a not applicable as EM images of LN have not been published previously.
<sup>N</sup> Lewy neurite
⊙ Difficult to distinguish filaments from clumped proteinaceous matter

**Figure S1.**
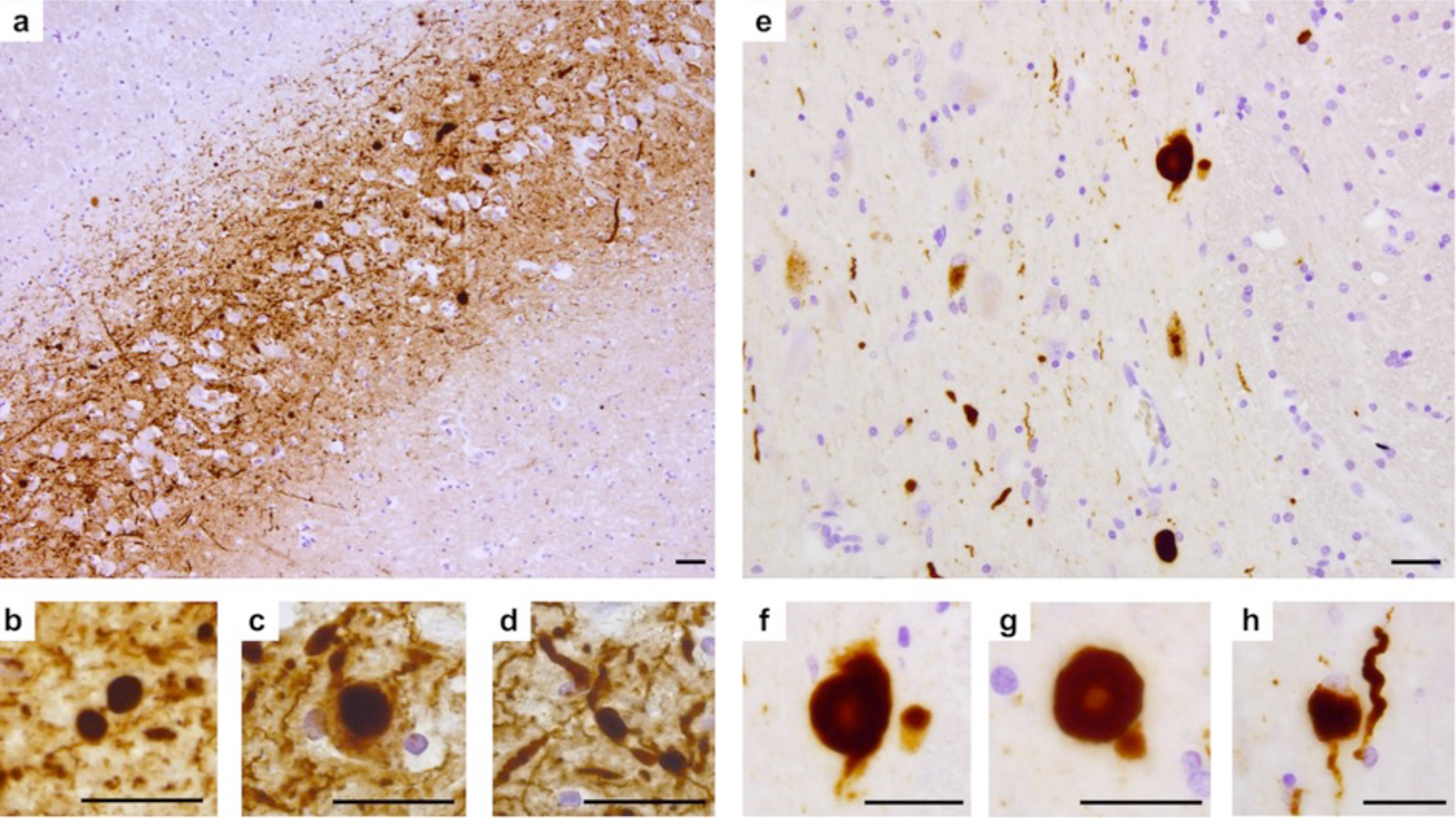
Immunohistochemical analysis of aSyn pathology. (**a-d**) aSyn (KM-51) immunostaining in the CA2 region of the hippocampus of Donor A-PD. (**e-h**) aSyn (KM-51) immunostaining of the *substantia nigra* of Donor B-PD. Images shown are from tissues that were taken from the same region of the same brain donors used for the other methods employed in this study, including CLEM and SBFSEM. Scale bars = 50 µm.

**Figure S2.**
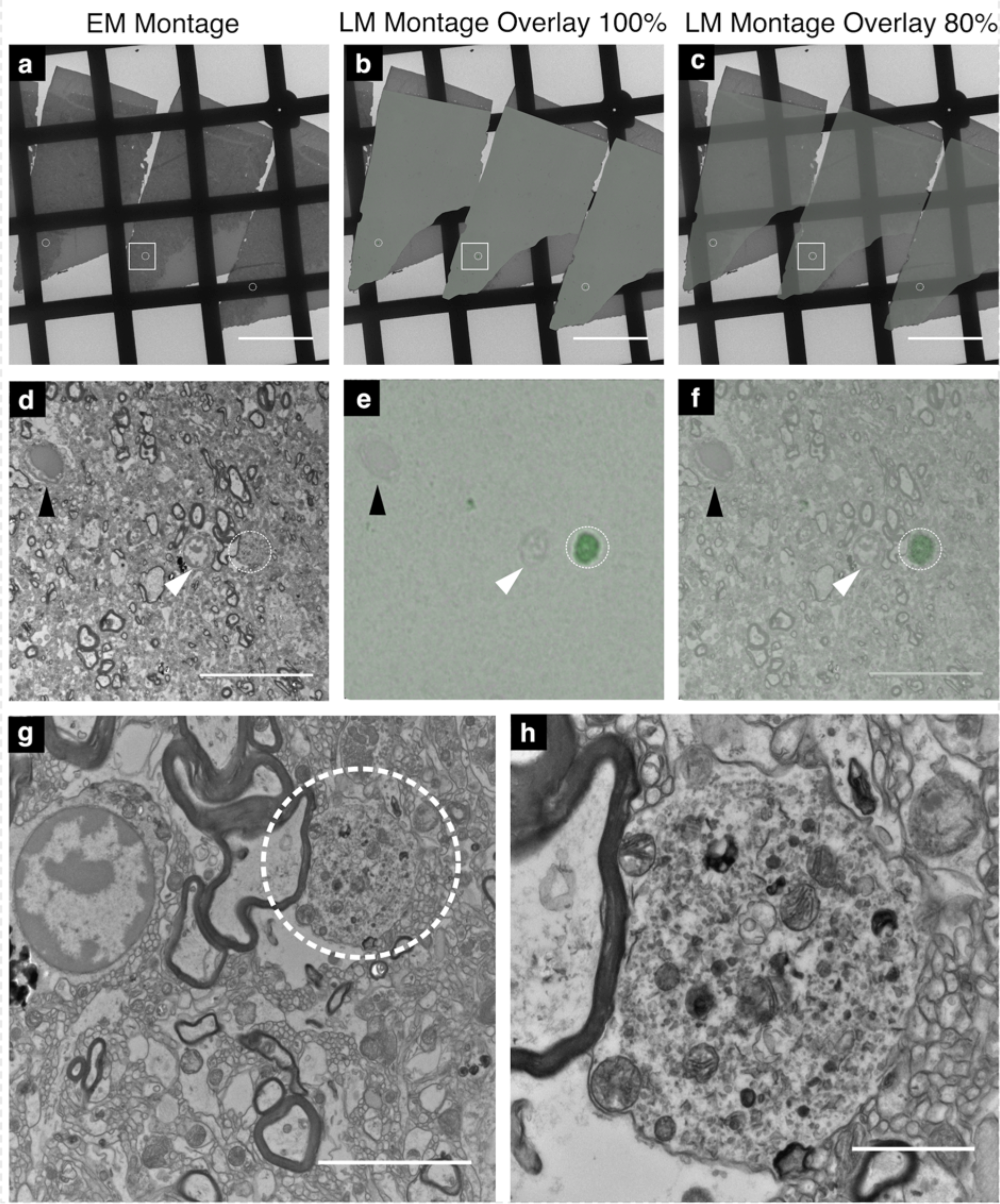
Correlative light and electron microscopy (CLEM) to identify Lewy pathology. aSyn-immunopositive inclusion from Donor D-PD is shown as an example. The same one inclusion serially sectioned is shown in each white circle in a-g. The same CLEM procedure shown here was applied to identify all Lewy structures in this study. (**a**) EM montage of 100-200 nm-thick tissue sections collected on an EM grid. **(b)** Light microscopy montage of aSyn-immunostained adjacent tissue sections (also 100-200nm-thick), overlaid onto the EM montage at 100% opacity. (**c**) Light microscopy montage overlay at 80% opacity. **(d-f)** Higher magnification area of the white box depicted in ‘a-c’; black arrowhead indicates blood vessel and white arrowhead indicates nucleus of nearby cell. Dotted white circle shows aSyn-immunopositive inclusion. **(e, f)** Colored feature represents inclusion, immunostained for aSyn; bound antibody complex detected by Permanent HRP Green Kit (Zytomed Systems), slides were counterstained with hematoxylin. **(g)** Higher magnification area of the sub-region shown in ‘d’ containing the inclusion (dotted circle) and neighboring nucleus. **(h)** Higher magnification of inclusion. Scale bars a-c = 200 µm, d-f = 20 µm, g = 5 µm, h = 1 µm.

**Figure S3.**
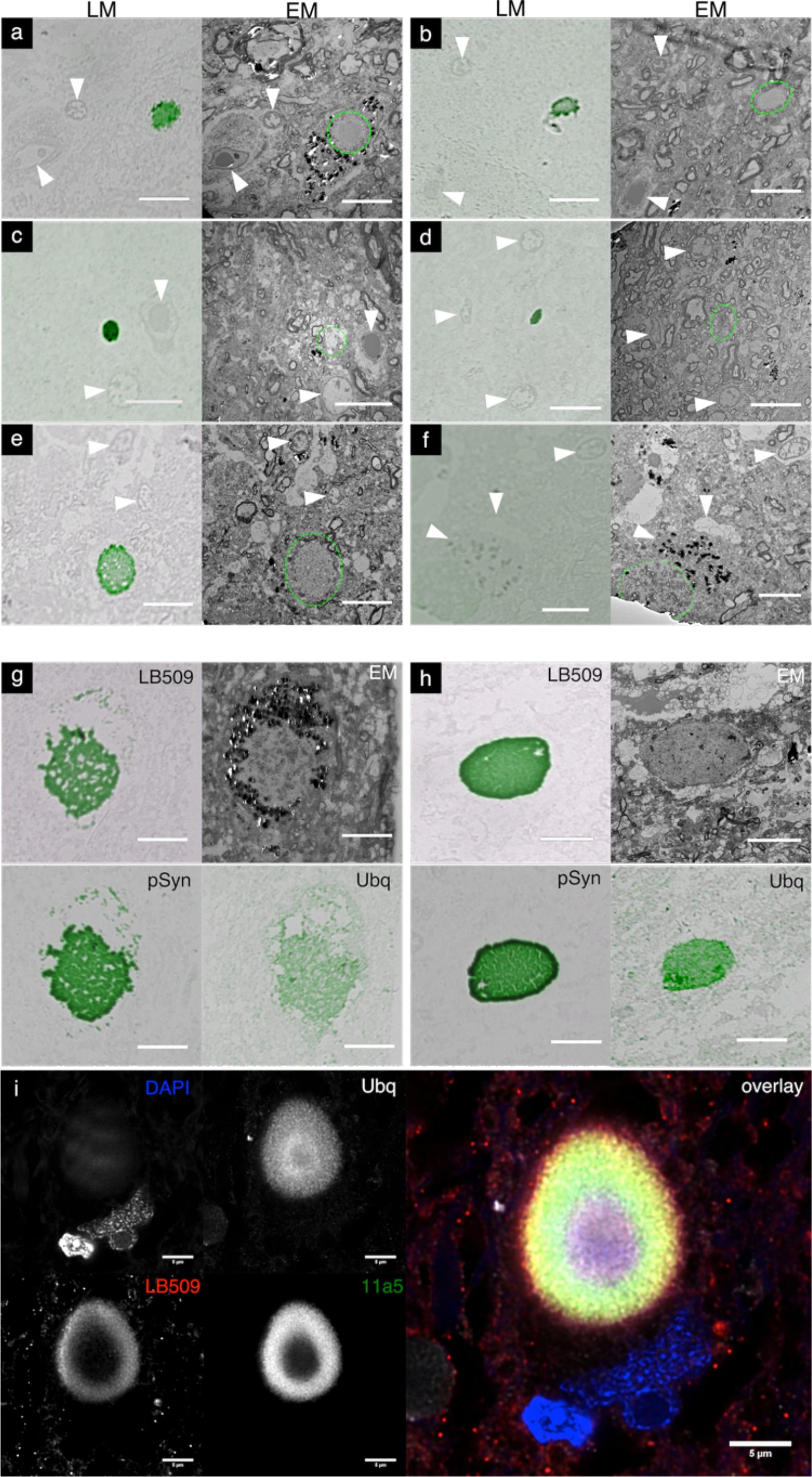
CLEM and CLSM to identify Lewy pathology. (a-f) Light microscopy (LM) image and correlating electron microscopy image (EM) for aSyn inclusions. 150 nm tissue sections collected on LM slides were processed using the LB509 antibody and immunopositive aggregates identified using a peroxidase detection system and green chromogen. Slides were counterstained with hematoxylin in order to identify cellular features for correlation with EM images. The same immunopositive inclusion is indicated (dashed green circle) in an adjacent 150 nm tissue section collected on an EM grid. White arrows indicate tissue features that were used for correlating the LM and EM images. (a) CLEM for Fig. 1d and S8, Donor C-PD, (b-d) CLEM for Fig. S6 d-f, respectively Donor D-PDD, (e-f) CLEM for Fig. S10 and S12, respectively Donor E-PD. (g,h) Examples of differential antibody staining for two aSyn immunopositive inclusions. Adjacent tissue sections were stained with either LB509, phosphorylated aSyn (pSyn; 11a5) antibody or ubiquitin (Ubq). The correlating EM picture for each inclusion is shown. (g) CLEM for Fig. S9, Donor D-PDD, (h) CLEM for Fig. S11, Donor D-PDD. All scale bars, a-h = 10 um. (i) CLSM images from a LB in a neuromelanin-containing neuron in the SN of Donor A-PDD, immunolabeled for alpha-synuclein (LB-509), Serine 129 phosphorylated alpha-synuclein (11A5) and ubiquitin (Ubq).

**Figure S4.**
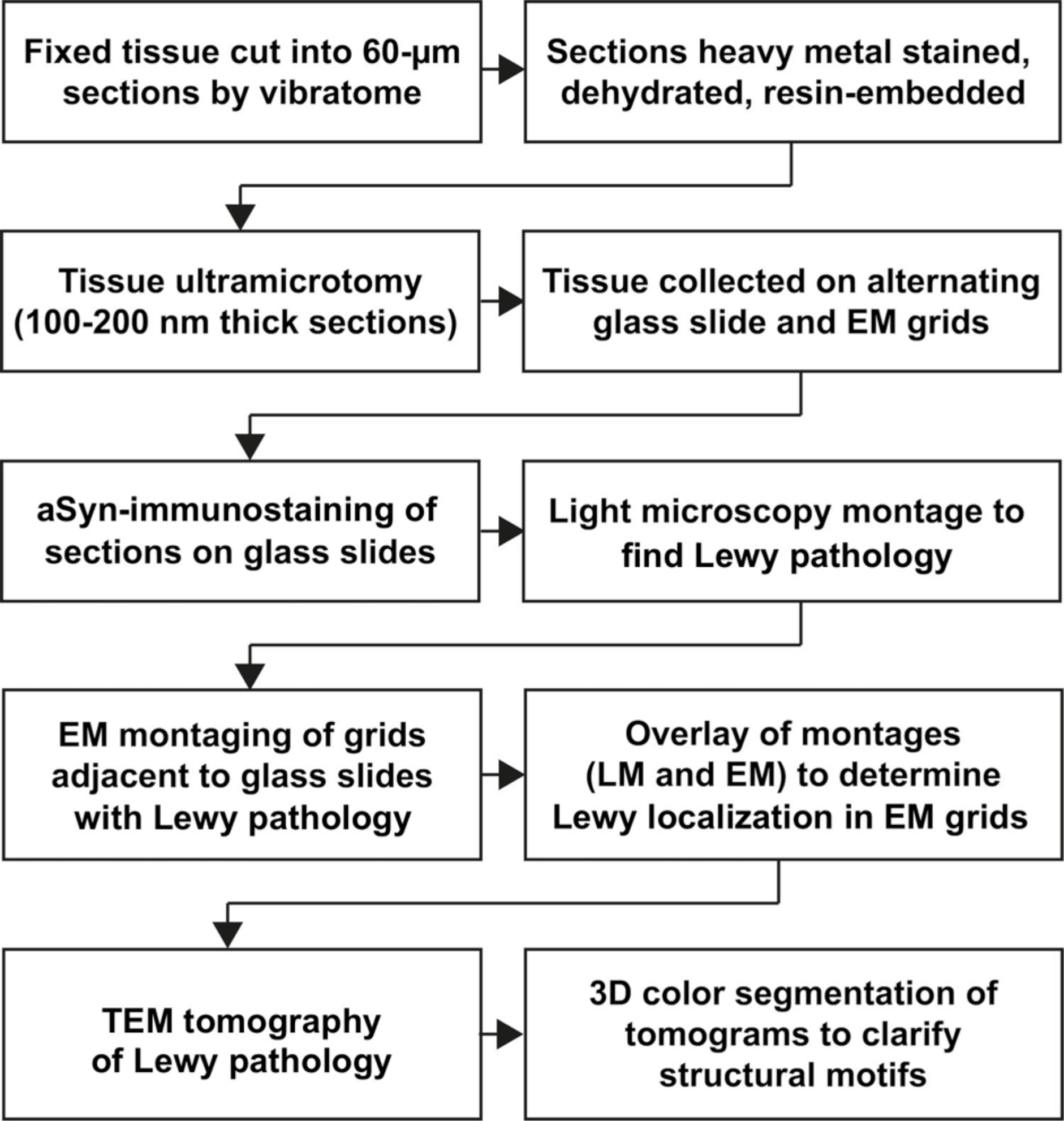
CLEM workflow. Correlative light and electron microscopy (CLEM) is often used to localize specific molecules of interest within the complex and diverse biological landscape of cells and tissues, typically via genetically encoded fluorescent or enzymatic markers^73^. Light microscopy is first used to visualize wide-field images with limited resolution, essentially providing a map to the labeled structures of interest. Such a map is then used to guide to the structure of interest for higher-resolution visualization by electron microscopy at a smaller imaging window. The general sequence of steps taken to achieve this for PD brain tissue sections is shown. EM = electron microscopy; LM = light microscopy.

**Figure S5.**
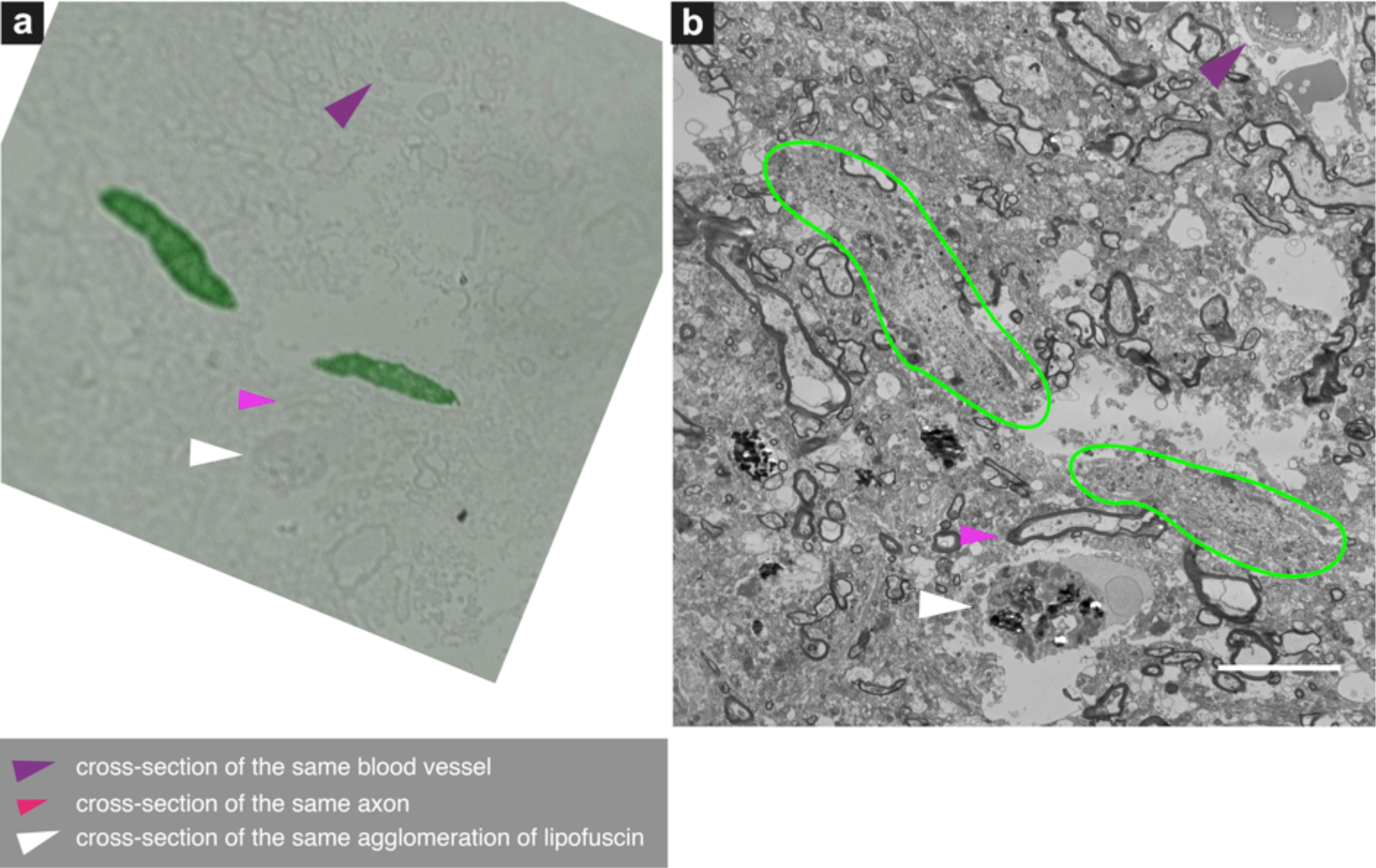
Correlative light and electron microscopy (CLEM) to identify Lewy neurites. aSyn-immunopositive inclusion from Donor E-PD is shown as an example. The essential procedure was used to identify all LN in this study. (a) Light microscopy image of aSyn-immunostained adjacent tissue sections (also 100-200nm-thick); Green colored features represents LN, immunostained for aSyn; bound antibody complex detected by Permanent HRP Green Kit (Zytomed Systems), slides were counterstained with hematoxylin. (b) Corresponding 2D EM image showing the same two regions of LN (circled in yellow) as identified by aSyn immunostaining in ‘a.’ Scale bar = 10 µm.

**Figure S6.**
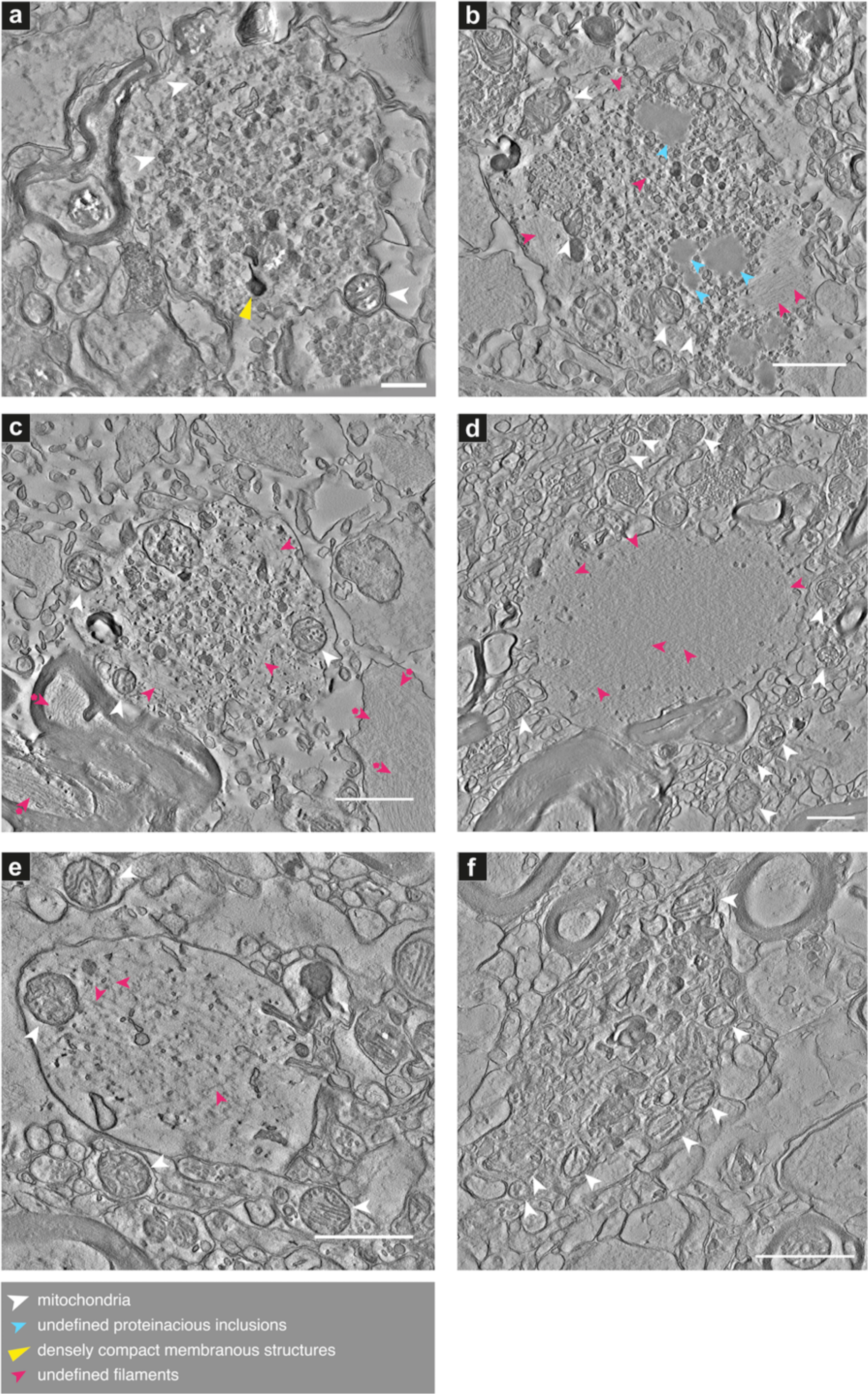
Lewy pathology as identified by CLEM. Projections of the central 20 slices of each reconstructed 3D tomogram are shown for each aSyn-immunopositive inclusion and surrounding cellular milieu. Feature details (arrowheads) are tabulated in Supplementary Table 1. Additional aSyn-immunopositive Lewy pathological inclusions are shown in Figs. 3, 4 and S5-S12. Donor identities are shown in Table 1. **(a)** aSyn-immunopositive inclusion in Donor A-PD (Movie 5), **(b-c)** in Donor B-PD (Movies 6, 7), **(d-f)** in Donor D-PD (Movies 8-10, CLEM data Fig. S3 b-d). Scale bars = 1 µm.

**Figure S7.**
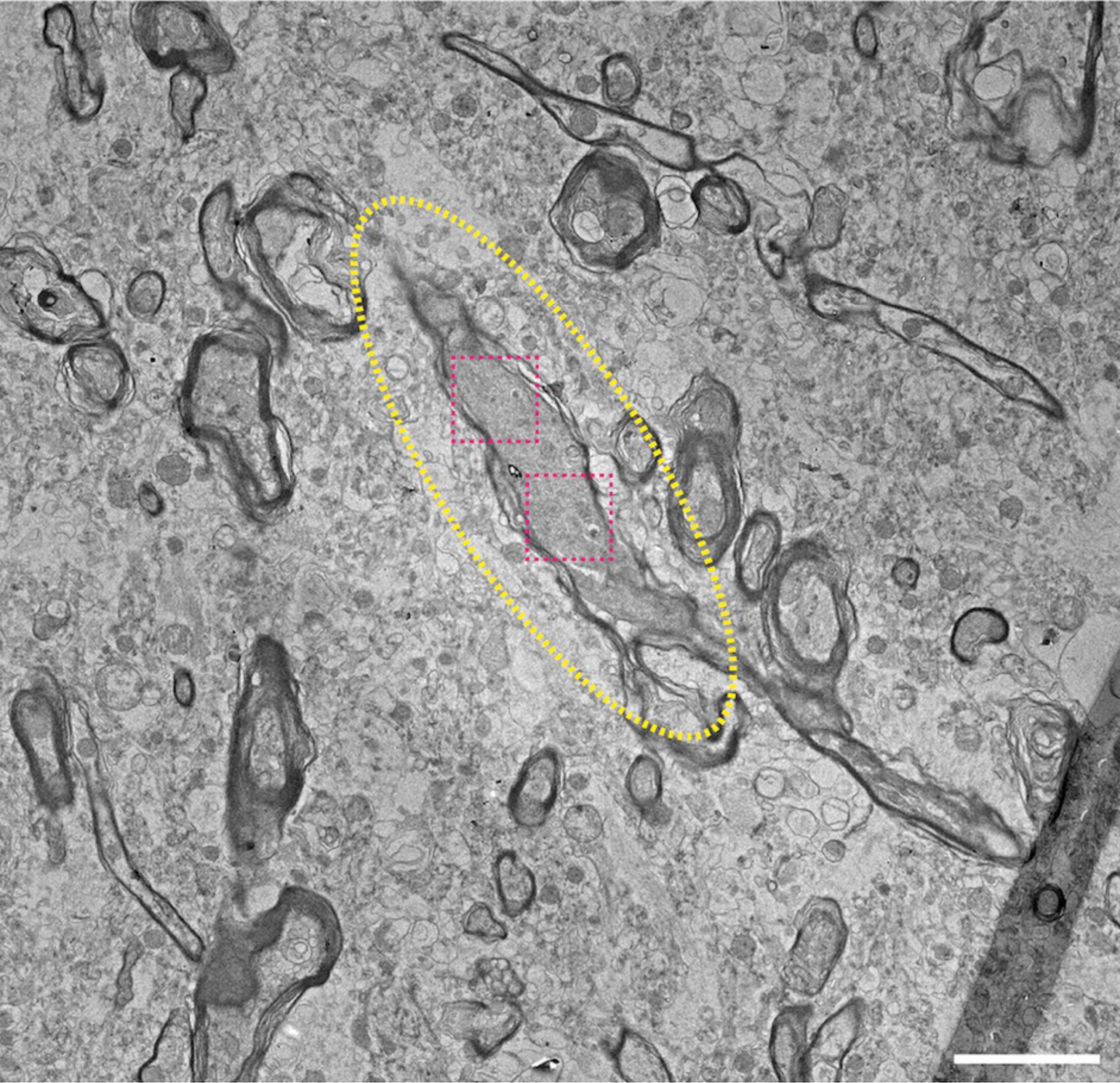
Low magnification overview of Lewy neurite identified using CLEM. From Donor B-PD, *substantia nigra*. 2D EM micrograph indicating the LN (yellow dotted oval) as identified by aSyn immunostaining in adjacent tissue section, and the specific positions where electron tomograms were collected (pink dotted boxes). Higher magnification images of pink dotted boxes represented in Figure 3. Scale bar = 3 µm.

**Figure S8.**
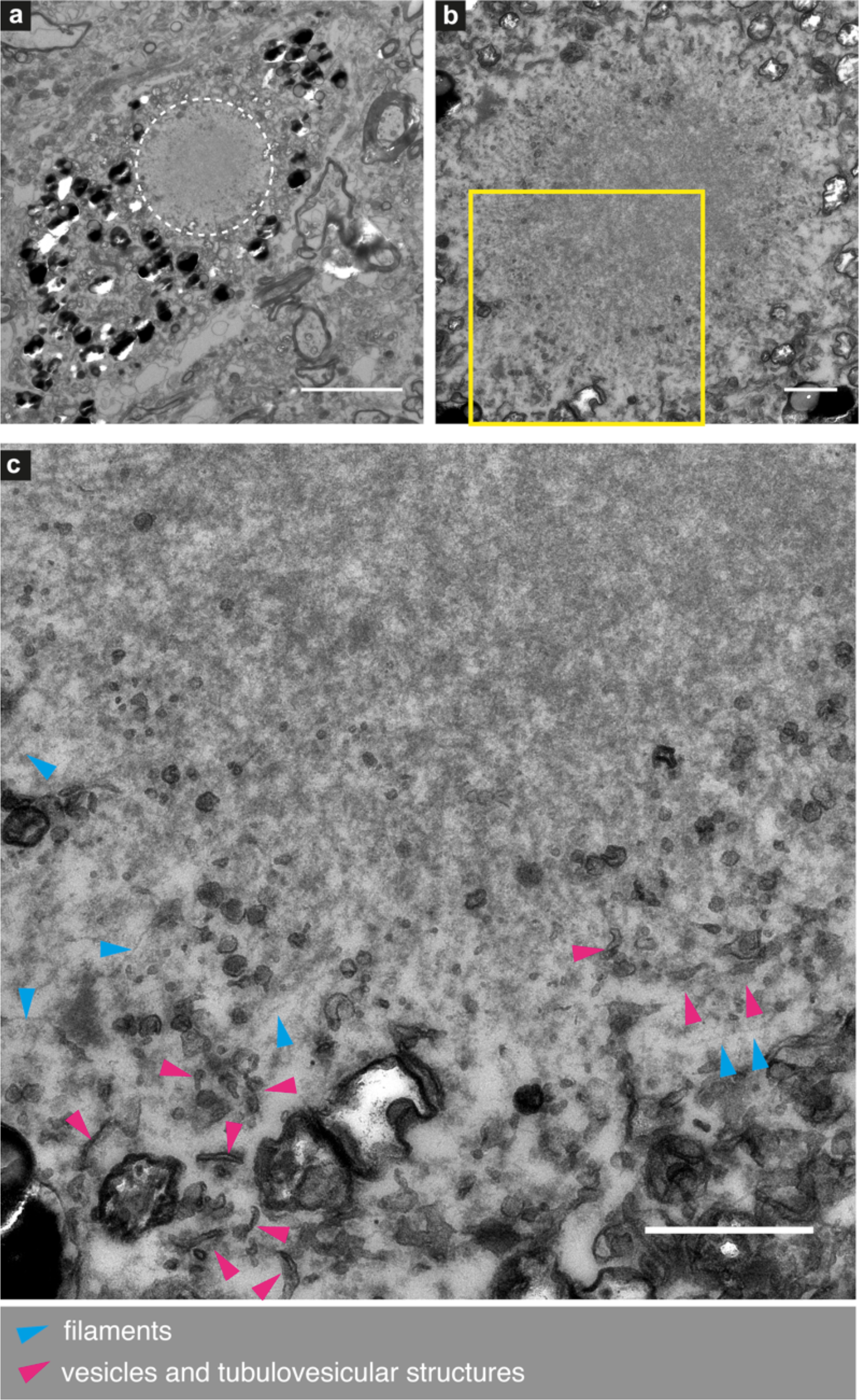
Filamentous Lewy pathology within neuromelanin-containing organelles. Identified by CLEM in Donor C-PD. CLEM data shown in Fig. S3a 2D electron micrographs showing the ultrastructure of a predominantly filamentous aSyn-immunopositive inclusion (same as shown in Fig. 1d) at **(a)** low magnification (white dotted circle) in which it can be seen amongst neuromelanin-containing organelles (black high contrast spots), and increasingly higher magnification in **(b)** and **(c)**. In addition to filaments and vesicles, distorted mitochondria are also interspersed at the periphery of the inclusion. Scale bars: a = 5 µm; b, c = 1 µm.

**Figure S9.**
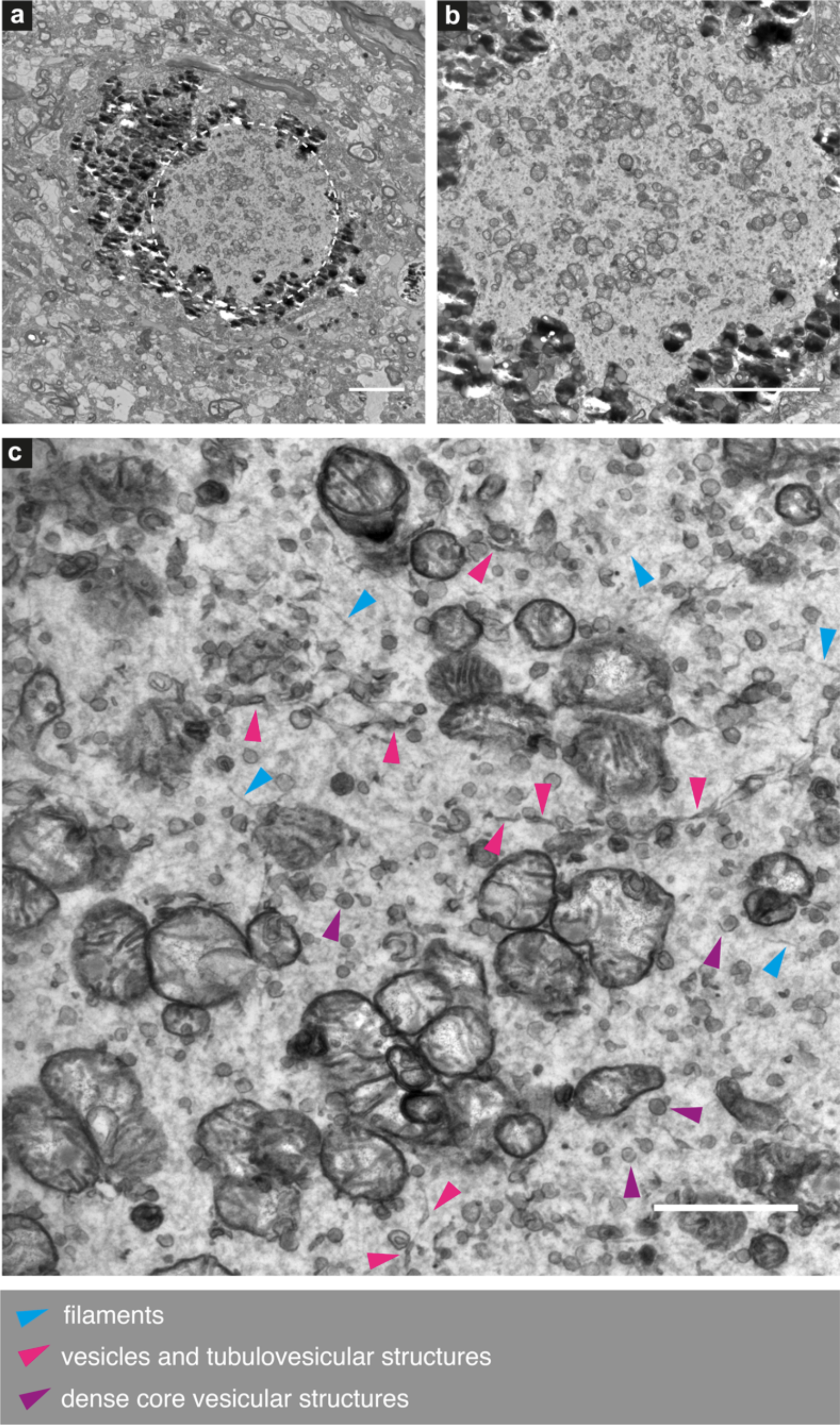
Membranous Lewy pathology within neuromelanin-containing organelles. Identified by CLEM in Donor D-PD. CLEM data shown in Fig. S3 g 2D electron micrographs showing the ultrastructure of a predominantly membranous aSyn-immunopositive inclusion at (a) low magnification (white dotted circle) in which it can be seen amongst neuromelanin-containing organelles (black high contrast spots), and increasingly higher magnification in (b) and (c). Abundant clustered mitochondria (vesicles with cristae) are interspersed amongst the other notated features. Scale bars: a, b = 5 µm; c = 1 µm.

**Figure S10.**
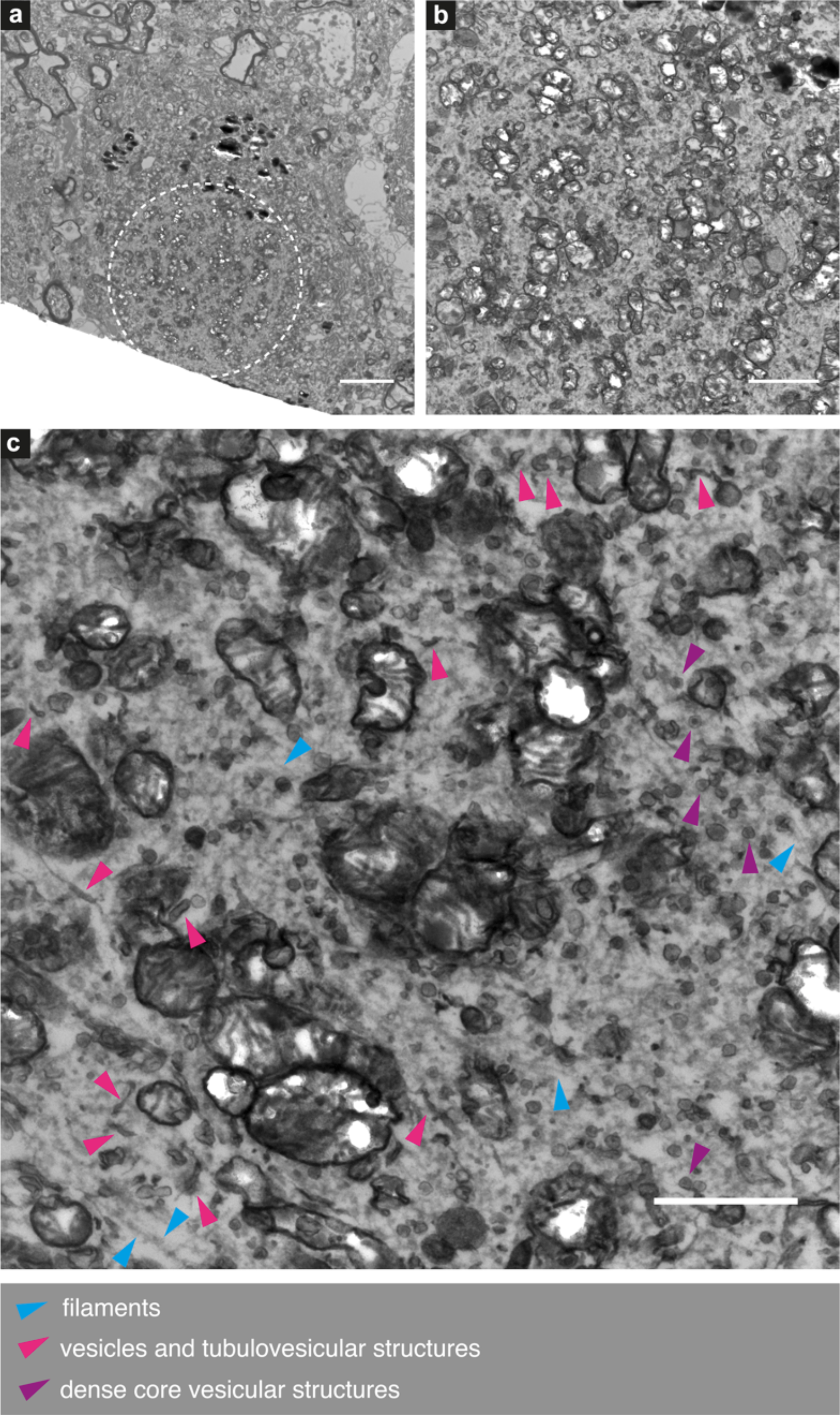
Membranous Lewy pathology within neuromelanin-containing organelles. Identified by CLEM in Donor E-PD. CLEM data shown in Fig. S3f 2D electron micrographs showing the ultrastructure of a predominantly membranous aSyn-immunopositive inclusion at **(a)** low magnification (white dotted circle) in which it can be seen amongst neuromelanin-containing organelles (black high contrast spots), and increasingly higher magnification in **(b)** and **(c)**. Abundant clustered mitochondria (vesicles with cristae) are interspersed amongst the other notated features. Scale bars: a = 5 µm; b = 2 µm; c = 1 µm.

**Figure S11.**
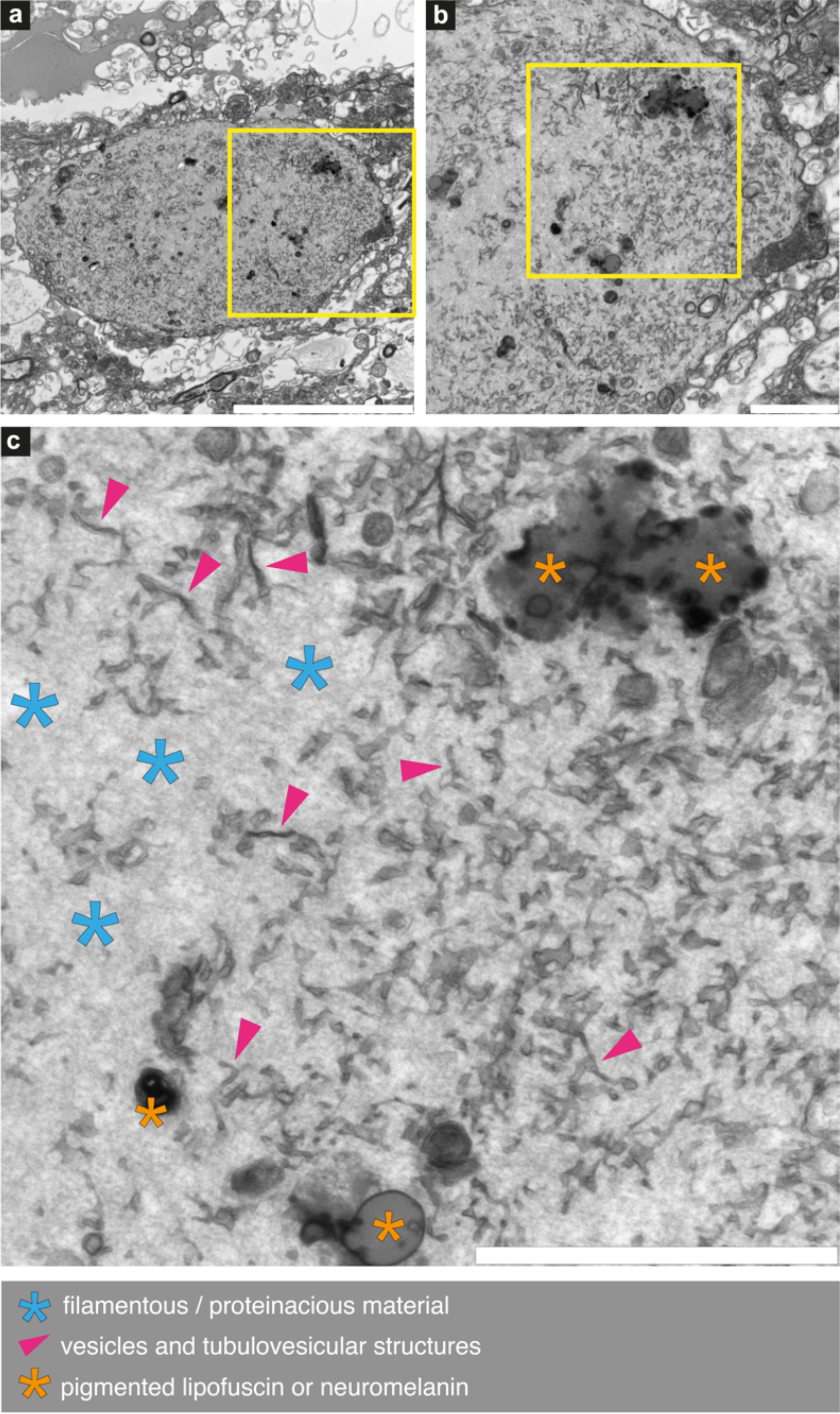
Lewy pathology consisting of abundant tubulovesicular structures. Identified by CLEM in Donor D-PD. CLEM data shown in Fig. S3h. 2D electron micrographs showing the ultrastructure of an aSyn-immunopositive inclusion at **(a)** low magnification in which it can be seen delimited by membrane, at increasingly higher magnification in **(b)** and **(c)** as indicated by the yellow boxes. Scale bars: a = 10 µm; b, c = 2 µm.

**Figure S12.**
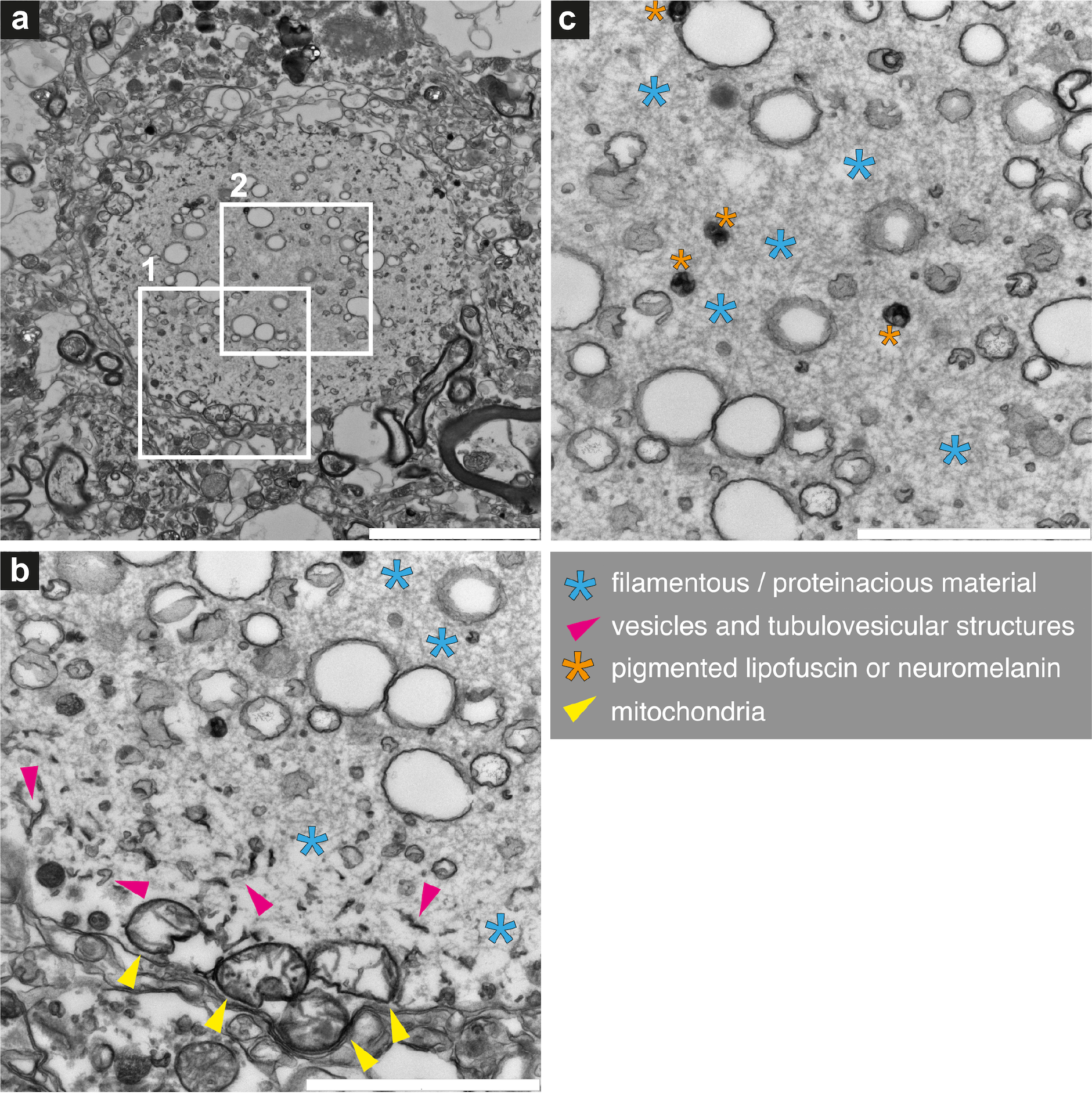
Lewy pathology consisting of abundant vesicular structures interspersed with filaments. Identified by CLEM in Donor E-PD. CLEM data shown in Fig. S3e 2D electron micrographs showing the ultrastructure of an aSyn-immunopositive inclusion at **(a)** low magnification and higher magnification of boxed region ‘1’ in **(b)** and ‘2’ in **(c)**. Abundant autophagic vacuolar-like structures (membrane-enclosed, “empty” vesicles) and vesicles with a ruffled border observed mainly in center. In addition to the other annotated features, abnormal mitochondria with few cristae are visible at the periphery (yellow arrowheads). Scale bars: a = 5 µm; b, c = 2 µm.

**Figure S13.**
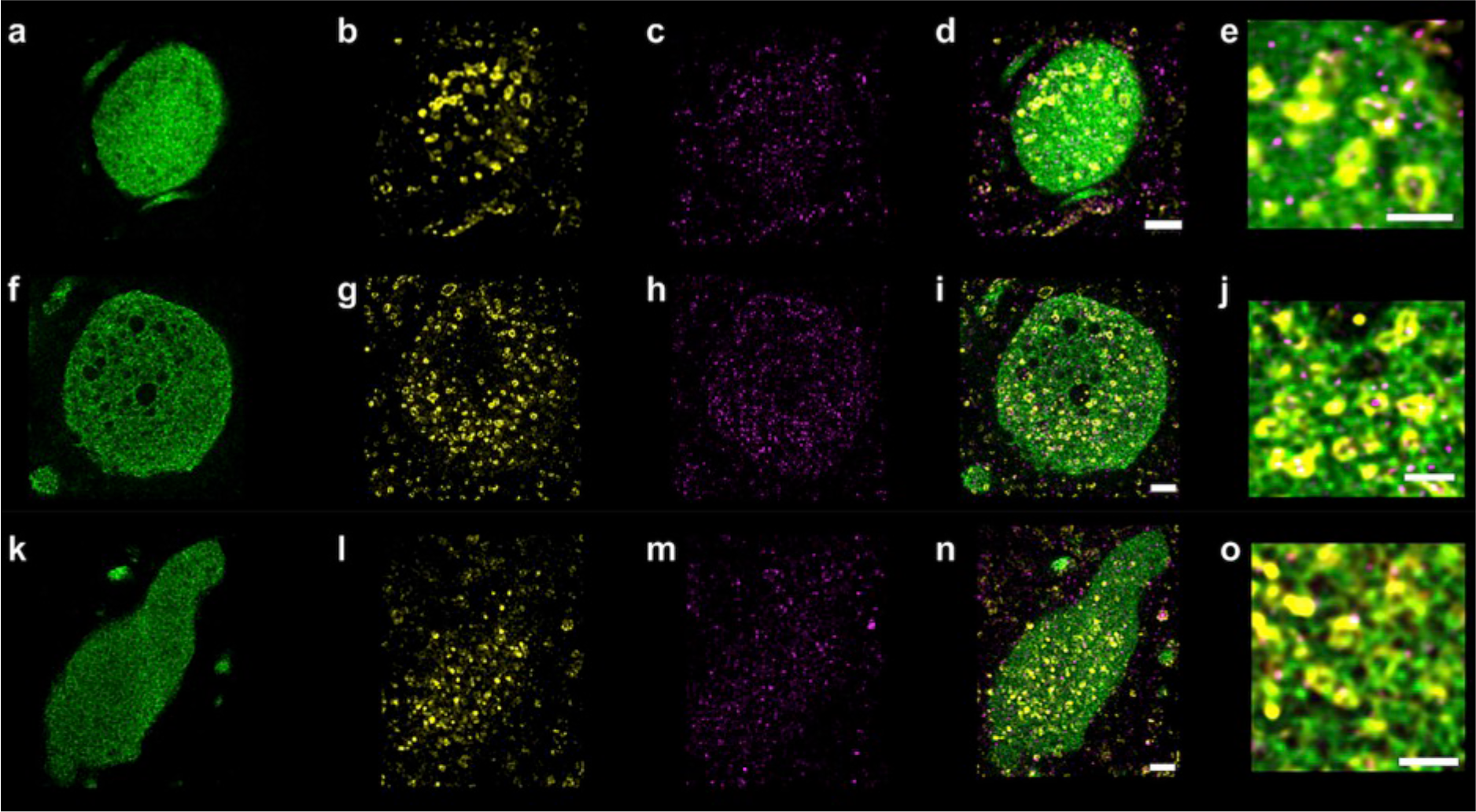
Subcellular distribution of aSyn and organelle markers within Lewy pathology without a p-aSyn positive outer layer. STED microscopy showing distribution of (a) marker for phosphorylated aSyn (pS129), (b) marker for mitochondria (porin VDAC1), (c) marker for lysosomes (LAMP1), (d) overlay of ‘a-c’, (e) higher magnification view of the edge of the aSyn-immunopositive inclusion shown in ‘d’ (f-i) Same STED microscopy and markers as in ‘a-d’, but a different inclusion, showing empty vacuoles that may represent autophagic vacuolar-like structures reminiscent of CLEM (Fig. S12), (j) higher magnification view of center of the inclusion shown in ‘i’. (k-n) Same STED microscopy and markers as in ‘a-d,’ but a LN, (o) higher magnification view of the LN as in ‘n.’. Images are representative across 14 PD donors for Lewy structures without the p-aSyn outer layer. Scale bars: d, i, n = 2 µm; e, j, o = 1 µm.

**Figure S14.**
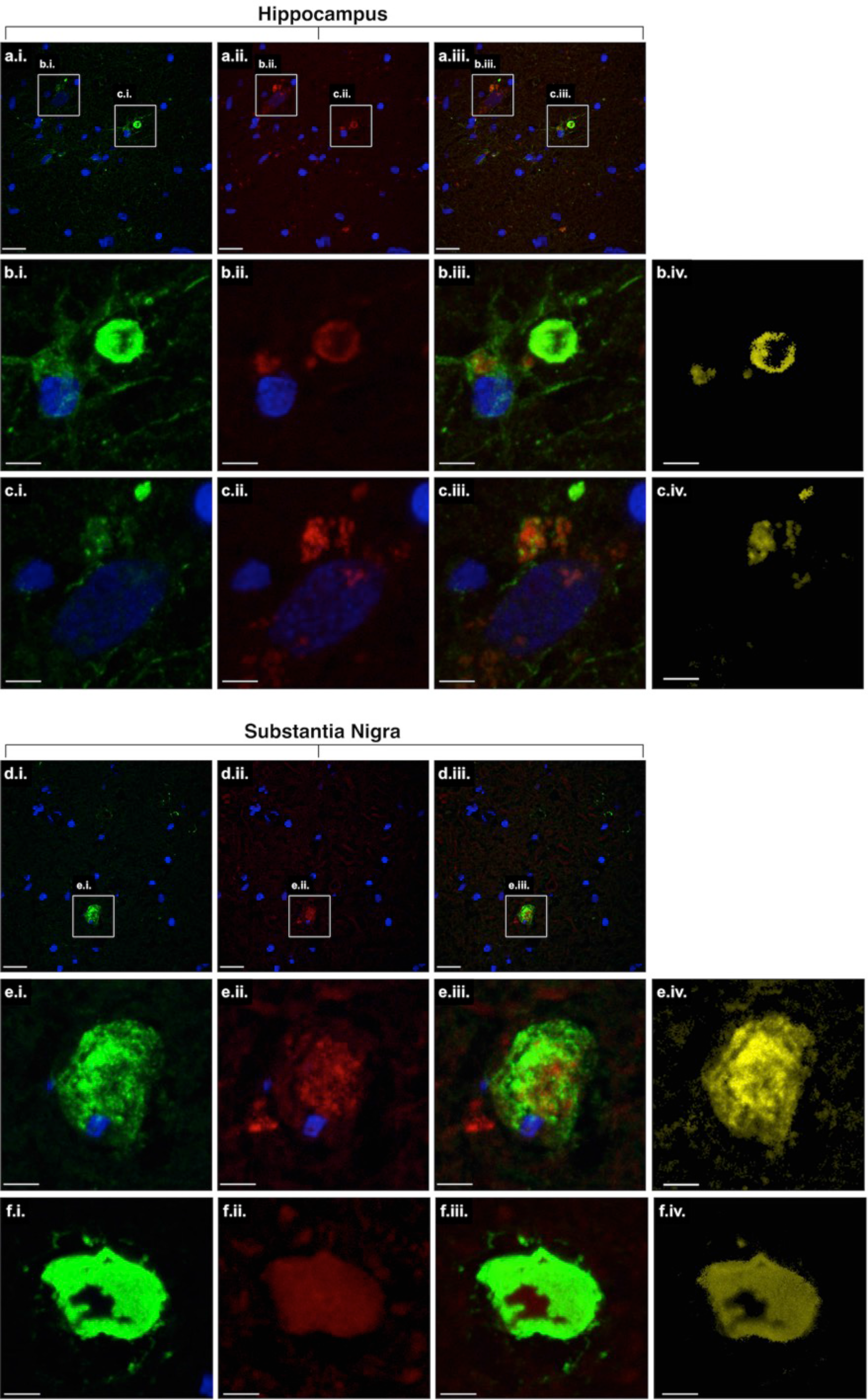
Co-localization of lipids with aSyn in Lewy pathology. Confocal fluorescence light microscopy projected image stacks of snap-frozen 10µm-thick tissue showing aSyn-immunopositive inclusions in the (**a-c**) hippocampal CA2 region of Donor A-PD, and (**d-f**) SN of Donor B-PD. Inclusions immunopositive for aSyn are visualized in green (LB509 antibody), lipid-rich structures are visualized in red by Nile Red staining, and nuclei are visualized in blue by DAPI. Column i = aSyn (green), nuclei (blue); Column ii = lipids (red), nuclei (blue); Column iii = overlay of aSyn (green), lipids (red), and nuclei (blue); Column iv = co-localization of aSyn and lipids (yellow). Scale bars: a, d = 20 µm; b, c, e, f = 5 µm.

**Figure S15.**
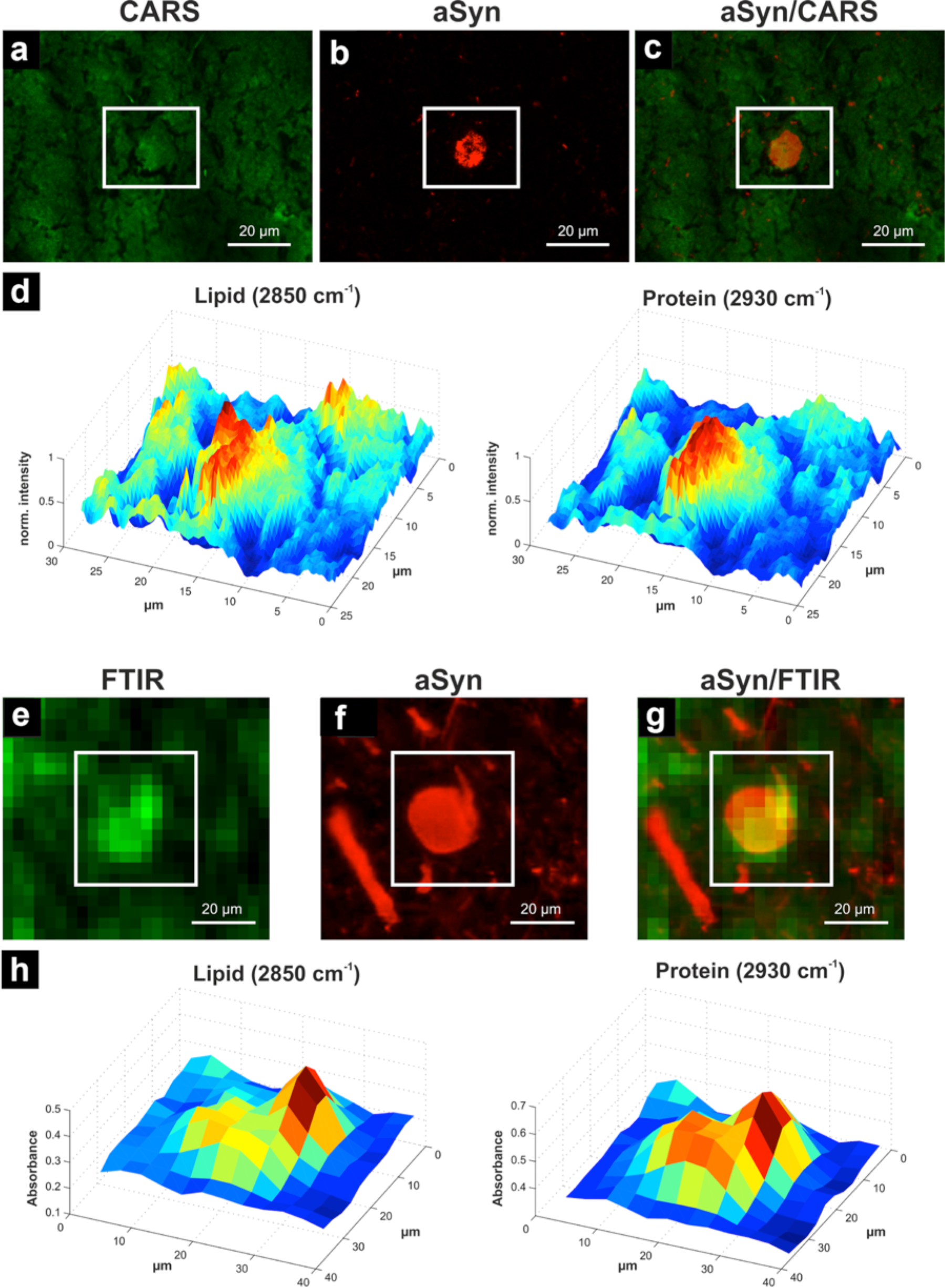
Lipid and protein distributions in Lewy pathology detected by label-free CARS or FTIR imaging combined with correlative immunofluorescence CLSM for aSyn. In Donor A-PD, CA2. (a) CARS image of lipids in an aSyn-immunopositive inclusion in PD brain tissue, recorded at 2850 cm^-1^. (b) Projected confocal immunofluorescence stack showing the same area, after immunostaining for aSyn (LB509). (c) Overlay of the CARS and aSyn immunofluorescence data shown in ‘a’ and ‘b’. (d) CARS intensity distribution profiles for lipids and proteins within the area, showing high peaks in the region of the LB. (e) FTIR image of lipids in an aSyn-immunopositive inclusion in PD brain tissue. (f) Projected confocal immunofluorescence stack showing the same area, immunostained for aSyn (LB509). (g) Overlay of the FTIR and aSyn immunofluorescence data shown in e and f. (h) FTIR intensity distribution profiles of lipids and proteins within the inclusion. Scale bars: 20 µm.

**Figure S16.**
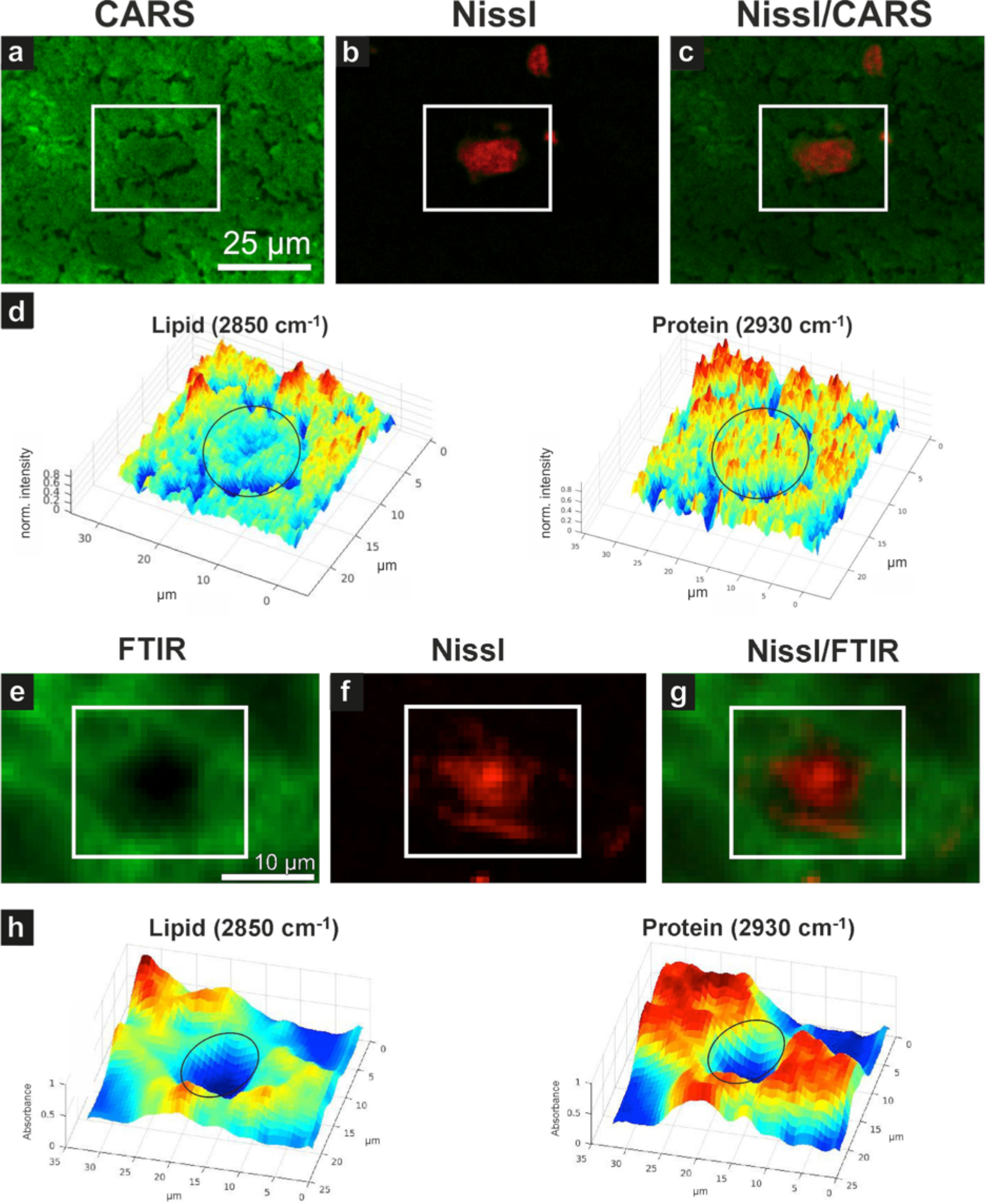
Detection of the lipid and protein distribution in neuron of control patient by label-free CARS and FTIR. (a) CARS image of lipids in a neuron of brain tissue, recorded at 2850 cm^-1^. **(b)** Confocal fluorescence showing the same area, after staining with Neurotrace (530/615) to stain the Nissl substance. **(c)** Overlay of the CARS and Neurotrace fluorescence data shown in ‘a’ and ‘b’. **(d)** CARS intensity distribution profiles for lipids and proteins within the neuron (black circle). These results show decreased lipid and similar protein intensities in neurons compared to neighboring tissue, which is in contrast with the results, i.e. increased lipids and proteins in Lewy structures (Figure S15). (**e**) FTIR image of lipids in a neuron of brain tissue, recorded at 2850 cm^-1^. **(f)** Confocal fluorescence showing the same area, after staining with Neurotrace. **(g)** Overlay of the CARS and Neurotrace fluorescence data shown in ‘e’ and ‘f’. **(h)** FTIR intensity distribution profiles for lipids and proteins within the neuron (black circle). These results show decreased lipid and similar protein intensities in neurons compared to neighboring tissue, which is in contrast with the results, i.e. increased lipids and proteins in Lewy structures (Figure S15).

**Figure S17.**
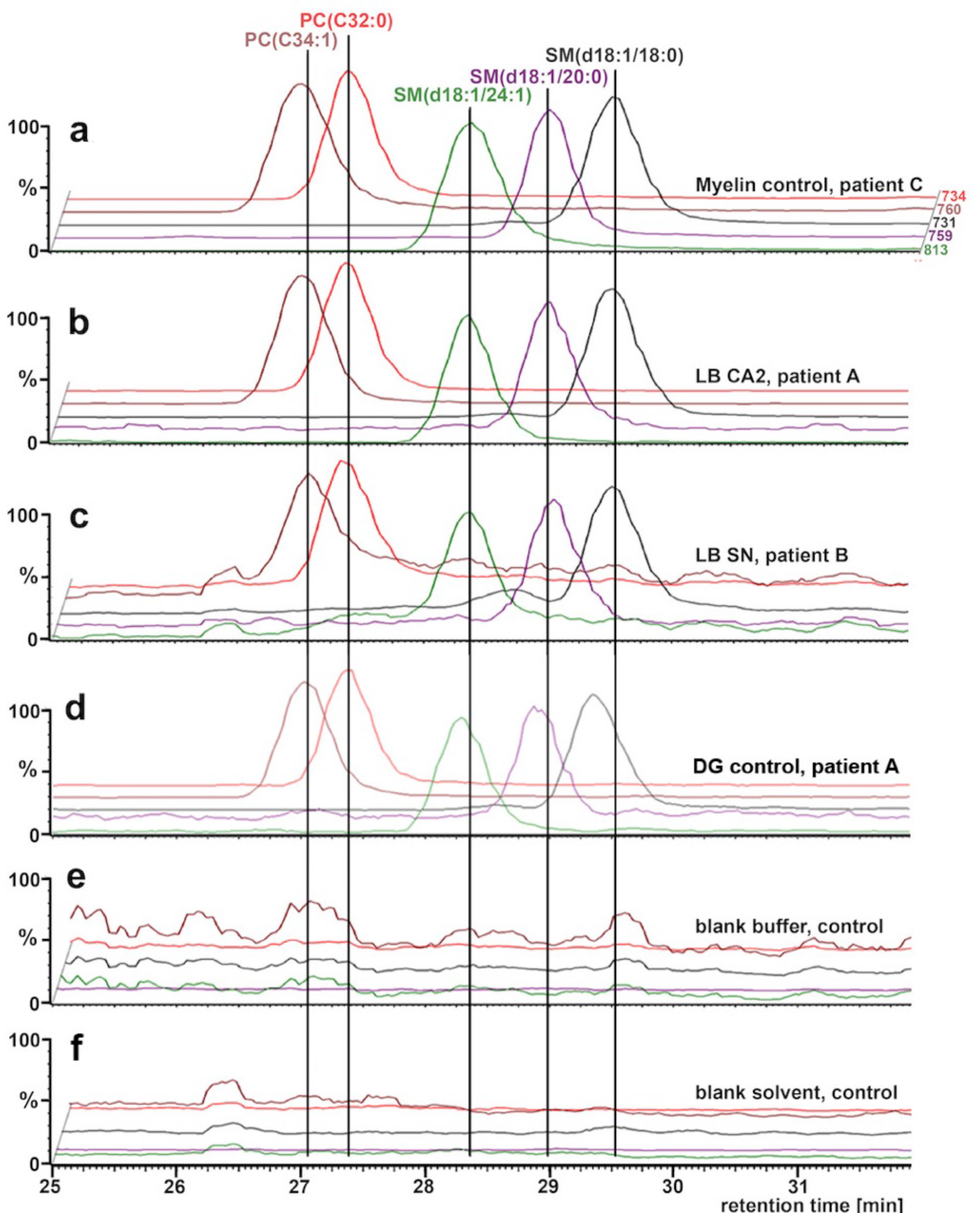
Liquid chromatography (LC) mass spectrometry (MS) and lipidomics reveal lipid content of Lewy pathology. Predominant peaks in all traces (**a–c**) represent the presence of phosphatidylcholine (PC) and sphingomyelin (SM) lipids. Mass spectrometric trace of (**a**) myelin as dissected from corpus callosum of non-neurological control donor, Donor F-Control, (**b**) laser capture micro-dissected Lewy bodies (∼2700) from hippocampal CA2 region of Donor A-PD, (**c**) laser capture micro-dissected Lewy bodies (∼3050) from *substantia nigra* of Donor B-PD. (**d**) Mass spectrometric traces of controls: dentate gyrus (DG) laser capture micro-dissected, not shown to contain any LB, from hippocampus of Donor A-PD; **(e)** blank tube, and (**f**) blank solvent, 5 µl chloroform/MeOH (2:1), as control experiments. Each curve shows MS signal over LC retention time with the m/z window scanning for the ratios indicated at the right end of the curves in ‘a’. These m/z windows select for the lipids indicated above the vertical lines in ‘a’.

**Figure S18.**
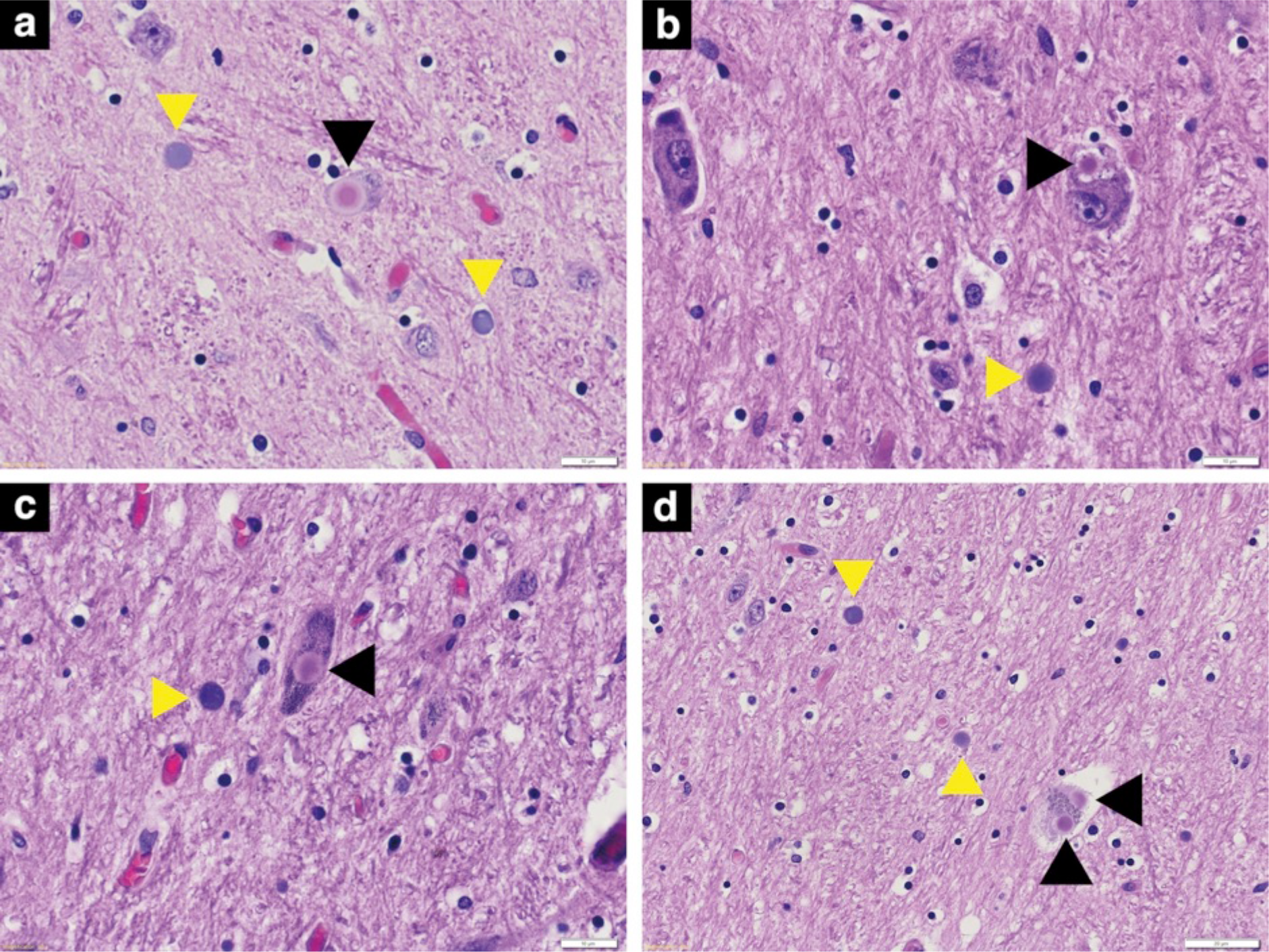
Conventional histopathological aspect of FFPE sample of the *substantia nigra* obtained from a PD brain donor shows Lewy pathology and *Corpora amylacea* side by side. (a-d): H&E stained tissue sections; yellow arrowheads indicate CA; black arrowheads indicate Lewy bodies. Note the similar size of the two structures and that they can occur in close proximity to one another. Scale bars: a-c = 10 µm; d = 20 µm.

## Movies

Movies are available on YouTube, at https://www.youtube.com/playlist?list=PLteVxMX-rlQu3HIZtObzPRspOr9uvA4s9 and individually at the links provided below.

**Movie 1:** Reconstructed and color-segmented 3D transmission electron tomogram of aSyn-immunopositive inclusion (LB). Corresponds to Fig. 1a. Thickness of tissue section imaged ≈ 150 nm.

https://youtu.be/StUx7Gxp3tI

**Movie 2:** Reconstructed and color-segmented 3D transmission electron tomogram of aSyn-immunopositive inclusion (LB). Corresponds to Fig. 1b. Thickness of tissue section imaged ≈ 150 nm.

https://youtu.be/FwZX3X6QkjQ

**Movie 3:** Reconstructed and color-segmented 3D transmission electron tomogram of aSyn-immunopositive inclusion (LB). Corresponds to Fig. 1c. Thickness of tissue section imaged ≈ 150 nm.

https://youtu.be/QrX73sGSbt4

**Movie 4:** Reconstructed and color-segmented 3D transmission electron tomogram of aSyn-immunopositive inclusion (LB). Corresponds to Figs. 1d, S5. Thickness of tissue section imaged ≈ 150 nm.

https://youtu.be/ROQ6mbj8VIA

**Movie 5:** Reconstructed and color-segmented 3D transmission electron tomogram of aSyn-immunopositive inclusion (LB). Corresponds to Fig. S4a. Thickness of tissue section imaged ≈ 150 nm.

https://youtu.be/7-2CN-V7NKQ

**Movie 6:** Reconstructed and color-segmented 3D transmission electron tomogram of aSyn-immunopositive inclusion (LB). Corresponds to Fig. S4b. Thickness of tissue section imaged ≈ 150 nm.

https://youtu.be/c82B7DCBIRE

**Movie 7:** Reconstructed and color-segmented 3D transmission electron tomogram of aSyn-immunopositive inclusion (LB). Corresponds to Fig. S4c. Thickness of tissue section imaged ≈ 150 nm.

https://youtu.be/ec5BG-ltlxM

**Movie 8:** Reconstructed and color-segmented 3D transmission electron tomogram of aSyn-immunopositive inclusion (LB). Corresponds to Fig. S4d. Thickness of tissue section imaged ≈ 150 nm.

https://youtu.be/iR6985Mp8qc

**Movie 9:** Reconstructed and color-segmented 3D transmission electron tomogram of aSyn-immunopositive inclusion (LB). Corresponds to Fig. S4e. Thickness of tissue section imaged ≈ 150 nm.

https://youtu.be/wg7v7I_BGsQ

**Movie 10:** Reconstructed and color-segmented 3D transmission electron tomogram of aSyn-immunopositive inclusion in neurite (LN). Corresponds to Fig. S4f. Thickness of tissue section imaged ≈ 150 nm.

https://youtu.be/Y0FaBmbWvpQ

**Movie 11:** Reconstructed and color-segmented 3D transmission electron tomogram of a region inside aSyn-immunopositive inclusion (LB, Fig. 1a) collected at higher magnification. Thickness of tissue section imaged ≈ 150 nm.

https://youtu.be/IwV0-xPiH5I

**Movie 12:** Reconstructed and color-segmented 3D transmission electron tomogram of a region inside aSyn-immunopositive inclusion (LB, Fig. 1a) collected at higher magnification. Tailed membrane stacks are clearly visible, as indicated in Fig. 1a (two yellow arrow-heads on right-hand side).

Thickness of tissue section imaged ≈ 150 nm.

https://youtu.be/9SDeEs5yJdQ

**Movie 13:** Reconstructed and color-segmented 3D transmission electron tomogram of region at the edge of the aSyn-immunopositive inclusion (LB, Fig. 1a) collected at higher magnification. A mitochondrion is clearly visible, as indicated in Fig. 2c (white oval). Thickness of tissue section imaged ≈ 150 nm.

https://youtu.be/2TwAJcmoH1g

**Movie 14:** Reconstructed and color-segmented 3D transmission electron tomogram of region at the edge of the aSyn-immunopositive inclusion (LB, Fig. S4a) collected at higher magnification.

Thickness of tissue section imaged ≈ 150 nm.

https://youtu.be/sgij8doJNzQ

**Movie 15:** Reconstructed and color-segmented 3D transmission electron tomogram of region inside the aSyn-immunopositive inclusion (LB, Fig. S4a) collected at higher magnification. Cluster of vesicles in separate adjacent compartment to LB is visible as shown in Fig. 2d. Thickness of tissue section imaged ≈ 150 nm.

https://youtu.be/IfcvCX133OU

**Movie 16:** Reconstructed and color-segmented 3D transmission electron tomogram of region within an aSyn-immunopositive Lewy neurite (same as shown in Fig. 3a) collected at high magnification. Thickness of tissue section imaged ≈ 150 nm.

https://youtu.be/ogdMLPaz_T0

**Movie 17:** Reconstructed and color-segmented 3D transmission electron tomogram of region within an aSyn-immunopositive Lewy neurite (same as shown in Fig. 3b) collected at high magnification. Thickness of tissue section imaged ≈ 150 nm.

https://youtu.be/D4r2PjtVy80

**Movie 18:** Reconstructed and color-segmented 3D transmission electron tomogram of region within a ‘control’ neurite in brain tissue from a non-demented, age-matched donor (same as shown in Fig. 3c) collected at high magnification. Thickness of tissue section imaged ≈ 150 nm.

https://youtu.be/Yln15OccuGs

**Movie 19:** Reconstructed and color-segmented 3D transmission electron tomogram of region within a ‘control’ neurite in brain tissue from a non-demented, age-matched donor (same as shown in Fig. 3d) collected at high magnification. Thickness of tissue section imaged ≈ 150 nm.

https://youtu.be/arL5GfyFgWM

**Movie 20:** Reconstructed serial block-face scanning electron tomograms depicting three separate Lewy pathological inclusions within the *substantia nigra* of Donor B. Scale bar = 5 µm.

https://youtu.be/O1Xb2LaELMI

**Movie 21:** Stimulated emission depletion microscopy showing a Lewy pathological inclusion in the same tissues (Donor B, *substantia nigra*) as taken from parallel blocks for the SBFSEM ultrastructural analysis (Fig. 3d). Thickness of tissue section = 20 µm.

https://youtu.be/dKO9HZqGTTI

